# TRF2 Non-Telomeric Function is Indispensable for Neural stemness

**DOI:** 10.1101/2025.02.28.640739

**Authors:** Soujanya Vinayagamurthy, Amit Kumar Bhatt, Sulochana Bagri, Supratim Ghosh, Arpan Parichha, Arindam Maitra, Shantanu Chowdhury

## Abstract

Depletion of TRF2 from chromosome ends results in telomeric fusions and genome instability in mammals. Here we show that although TRF2 is indispensable for the proliferation and survival of mouse neural stem cells (mNSCs), surprisingly, this is due to non-telomeric transcriptional function of TRF2, and not telomere protection. Complementing recent work showing TRF2 is dispensable for telomere protection in pluripotent stem cells. Deletion of TRF2 in adult mNSCs (TRF2^fl/fl^, Nestin-Cre) resulted in markedly reduced proliferation and impaired differentiation into neurons. However, telomere dysregulation-induced DNA damage was not observed, as indicated by the unaltered DNA damage response. Similarly, in SH-SY5Y cells, TRF2 depletion induced differentiation without causing telomere dysfunction. Mechanistically, non-telomeric TRF2 directly binds to the promoters of key genes that regulate differentiation. TRF2-dependent recruitment of the polycomb repressor complex (PRC2) and subsequent H3K27 trimethylation repress differentiation-associated genes, thereby maintaining NSC identity. Interestingly, G-quadruplex (G4) motifs are necessary for TRF2 binding. Disrupting the TRF2-G4 interaction— either through G4-binding ligands or the G4-specific helicase DHX36—induces differentiation genes, thereby promoting neurogenesis. These findings reveal a pivotal non-telomeric role of TRF2 in NSC survival, providing key mechanistic insights into neurogenesis with implications for aging-related neurodegeneration.

## Introduction

The primary function of TRF2 (Telomeric Repeat-binding Factor 2) is to protect telomeres. Telomere capping by TRF2, a component of shelterin, prevents telomeres from being detected as double-strand DNA breaks [1,2]. However, recent work from Denchi and Boulton groups showed TRF2-mediated telomere protection was dispensable in mouse embryonic stem cells (mESCs) [3,4]. These studies suggested TRF2’s telomeric function may vary depending on cell type or developmental stage.

Notably, TRF2 remains elevated in the brain throughout development and adulthood, in contrast to other tissues where TRF2 levels decline after embryonic day 16 [5]. Further, TRF2 is upregulated when human embryonic stem cells (hESCs) commit to the neural lineage and TRF2 levels are maintained in neural progenitor cells [6]. Deficiency in the TRF2 ortholog (*terfa*) in zebrafish show neurodevelopmental defects [7]; while in mice, TRF2 knockout resulted in telencephalic cell death during embryonic development [8]. Moreover, in differentiated neurons TRF2 was found to be primarily cytoplasmic, and present as the truncated form (TRF-S) lacking the DNA binding domain [9]; whereas in NSCs it is nuclear and reported to interact with the RE1 silencing factor REST [6]. Although, these underline the importance of TRF2 in NSCs, the role of TRF2 – telomeric and/or non-telomeric – in NSCs remains poorly understood.

Relatively recent studies, including from our group, show TRF2 functions as a transcriptional regulator by binding to non-telomeric promoter sites spread across the genome, in addition to telomere protection [10–13]. Non-telomeric TRF2 was found to recruit epigenetic complexes to induce permissive/closed chromatin leading to altered gene expression [14–16]. In multiple instances, interestingly, TRF2 binding to promoter DNA required presence of intact G-quadruplex (G4) structure(s) [14–18]. Consistent with extensive (>20000) non-telomeric TRF2 binding sites found genome wide from ChIPseq wherein a majority of the sites comprised potential G4 sequences [18].

Potential G4s have been reported to be enriched at promoters [19,20], and critical for transcription factor binding including epigenetic functions that are G4 dependent [11,15,21,22]. Contextually, notable are gene ontology analyses showing enrichment of promoter G4s in genes related to particularly neurogenesis [19,20]. With these, here we asked: (a) If TRF2 binds to promoters of genes involved in neuronal differentiation, and how this might impact cell fate of neural stem cells (NSC); and (b) if TRF2 is required for telomere protection in NSCs.

Our findings demonstrate TRF2 is critical for maintaining neural stemness, but not for telomere protection. TRF2 depletion did not induce telomeric damage and DNA damage response (DDR) was not activated. Non-telomeric TRF2 was found to bind directly to promoter DNA, and repress key neuronal differentiation genes. The repression was through TRF2-dependent recruitment of the RE-1 silencing factor (REST) and the canonical Polycomb Repressive Complex 2 (PRC2). Upon TRF2 loss, the neuronal differentiation genes were upregulated promoting neuronal differentiation. Together these underline the critical role of TRF2 in non-telomeric transcriptional regulation essential for maintaining the neural stem state, but dispensable for telomere protection in NSC.

### DNA binding by TRF2 is critical for maintaining the neural stem state

Using tissue from adult mouse brain we first checked for presence of TRF2 within the dentate gyrus, the adult neural stem cell niche, and other regions of the hippocampus (Fig S1.1 a). This confirmed presence of TRF2 within cells in the dentate gyrus. Next, we performed single cell RNA (scRNA-seq) of the adult mouse hippocampus. Using RNA expression from 4783 single cells from this experiment we compared TRF2 expression across different cell types and subtypes of neural stem cell (NSC) populations in the dentate gyrus (Fig S1.1 (b-h)). This interestingly showed the NSC subtype with high-SOX2 (although low in number as reported earlier) also had enhanced levels of TRF2 relative to low-SOX2 NSCs (Fig S1.1 (e-f)). This was consistent with results from analyses of publicly available spatial transcriptomic (Fig S1.2 a,b), single-nuclei (S1.2 c-g) and single-cell datasets from adult and embryonic mouse brains (Fig S1.3) which showed presence of TRF2 in SOX2-high NSCs [23–25]. Together this support presence of TRF2 within the adult NSC population. In addition, we noted TRF2 within the differentiated neurons (βIII-Tubulin-high); which is likely the truncated non-nuclear form reported earlier [5,9,26].

Next, we sought to understand the role of TRF2 in mouse Neural Stem Cells (mNSCs). The “floxed” mouse TRF2 allele (TRF2-flox) [27] was crossed with a strain expressing 4-hydorxy tamoxifen (4-OHT) inducible Cre recombinase under the control of the Nestin promoter (nestinCreERt2) [28]. This resulted in nestinCreERt2:TRF2-flox [conditional knockout (cKO)] transgenic line where 4-OHT treatment resulted in deletion of TRF2 in the nestin-expressing neural stem/progenitor cells (Fig 1a). Hippocampus was isolated from the brains of nestinCreERt2:Trf2-flox transgenic strain and TRF2 deletion was induced with 4-OHT in neurosphere culture. TRF2 deletion in NSCs resulted in significant reduction in number of neurospheres compared to vehicle control (EtOH) NSCs (Fig 1b). Reduction in Ki67 expression (Fig 1c, i), in addition to increase in MAP2, BDNF and L1CAM neuronal differentiation markers (1c, ii-1v) was clear. This indicates a reduction in the NSC population along with reduced proliferative capacity upon loss of TRF2.

**Figure 1:**
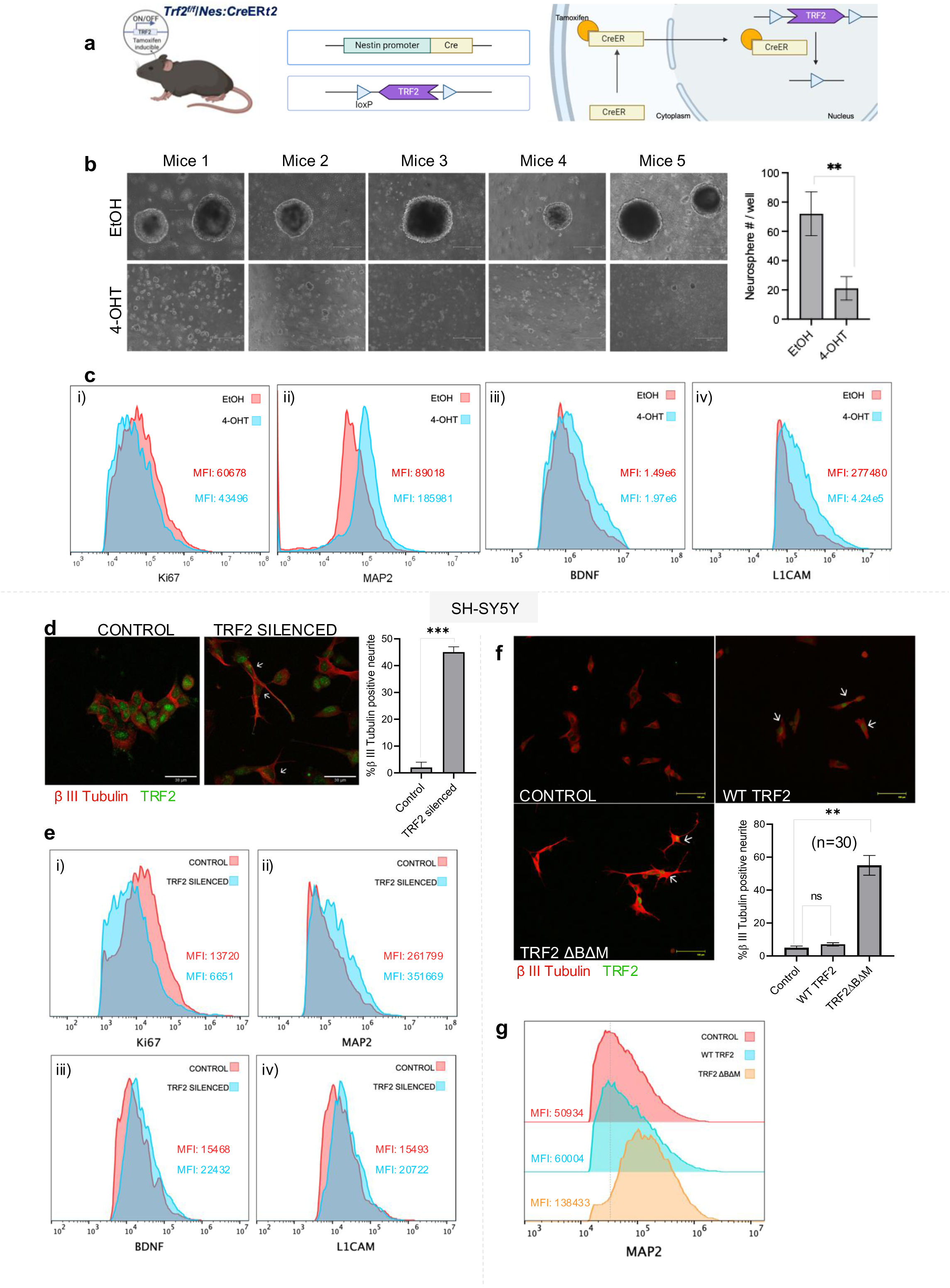
TRF2 inhibition triggers neuronal differentiation. a. Schematic of *Terf2^F/F^*/*Nes:Cre* mice. b. Neurosphere assay with mNSCs from *Terf2^f/f^*/*Nes:Cre* mice. 5 days post treatment with (4– OHT) tamoxifen and (EtOH) vehicle control. The graph indicates the number of neurospheres formed per well (error bar ± SD (unpaired t test)) c. Flow cytometry of mNSCs from *Terf2^F/F^*/*Nes:Cre* mice; Vehicle control (EtOH) and TRF2 knockout cells (4-OHT). Mean intensity of fluorescence (MIF) for (i) Ki67, (ii) MAP2 (iii) BDNF, (iv) L1CAM and TRF2 (supplementary) is shown. The cell counts are normalized to respective modes for comparative representation. d. Immunofluorescence staining of TRF2 and β-III Tubulin proteins in normal and TRF2 silenced SH-SY5Y cells. β- III Tubulin and TRF2 were stained using Alexa fluor-594 (red signal) and Alexa fluor-488 (green signal), respectively. The graph shows the quantification of β- III Tubulin positive neurite from 30 cells (n = 30) shown in respective right panels (error bars ± SE). e. Flow cytometry of Control and TRF2 silenced SH-SY5Y cells. Mean intensity of fluorescence (MIF) for (i) Ki67 (ii) MAP2, (iii) BDNF, (iv) L1CAM and TRF2 (supplementary) is shown. The cell counts are normalized to respective modes for comparative representation. f. Immunofluorescence staining for TRF2 and β- III Tubulin proteins in Control, WT TRF2 and TRF2 ΔBΔM overexpression conditions. β- III Tubulin and TRF2 were stained using Alexa fluor-594 (red signal) and Alexa fluor-498 (green signal), respectively. Quantification of β- III Tubulin positive neurite from 30 cells (n = 30) shown in respective right panels (error bars ± SE). g. Flow cytometry using staining for MAP2 and TRF2 in SH-SY5Y cells. Control, WT TRF2 and TRF2 ΔBΔM overexpression conditions. Mean intensity of fluorescence (MIF) for MAP2 and TRF2 is shown (supplementary). The cell counts are normalized to respective modes for comparative representation

We next investigated the effect of TRF2 on SH-SY5Y human neuroblastoma cell line. These cells originate from neural crest cells and exhibit key features of neuronal progenitors making them a widely used model to study neuronal differentiation [29–32]. Silencing TRF2 in SH-SY5Y cells resulted in enhanced neurite growth (Fig 1d). The decrease in Ki67 (Fig 1e, i), increase in MAP2 expression (Fig 1e, ii) along with BDNF, L1CAM (Fig 1e, iii,iv) was evident. Collectively, these results showed loss of TRF2 promotes neuronal differentiation.

Further we asked, whether the DNA binding activity of TRF2 is required for differentiation. Induction of the TRF2-DelB-DelM (TRF2ΔBΔM) mutant (lacking both the Basic and Myb domains of TRF2 required for DNA binding [33–35]promoted differentiation; whereas the full-length wild type (WT) TRF2 did not (Fig. 1m). Further, as expected, elevated MAP2 in case of TRF2ΔBΔM, and insignificant change in WT was observed (Fig 1l). Together, these support TRF2’s direct DNA binding to be key for maintaining stemness in NSCs.

### TRF2 mediated differentiation is independent of telomere damage

Next, we asked whether TRF2-loss affected telomere protection and resultant DNA damage in NSCs. On TRF2 knockdown following 4-OHT treatment in mNSCs the canonical DDR markers □H2AX, 53BP1, and p-ATM levels were not elevated compared to cells with vehicle control (Fig 2a). Similarly, in human SH-SY5Y cells, TRF2 loss did not result in enhanced levels of □H2AX, 53BP1, and p-ATM. Consistent with this, induction of TRF2ΔBΔM, devoid of DNA binding, in SHSY5Y cells did not show increase in the DDR markers (Fig 2c). Together these suggest lack of TRF2 did not induce telomere dysregulation in NSC. This was further tested by checking telomere dysfunction-induced 53BP1 foci at the telomeres [36]. Localization of 53BP1 to telomeres on TRF2 depletion in SH-SY5Y cells was insignificant compared untreated cells (Fig 2d). In contrast, as expected, TRF2 depletion resulted in significant increase in DDR markers (Sup Fig 2a) and telomere dysfunction-induced 53BP1 foci in somatic foreskin fibroblasts (Fig 2d).

**Figure 2:**
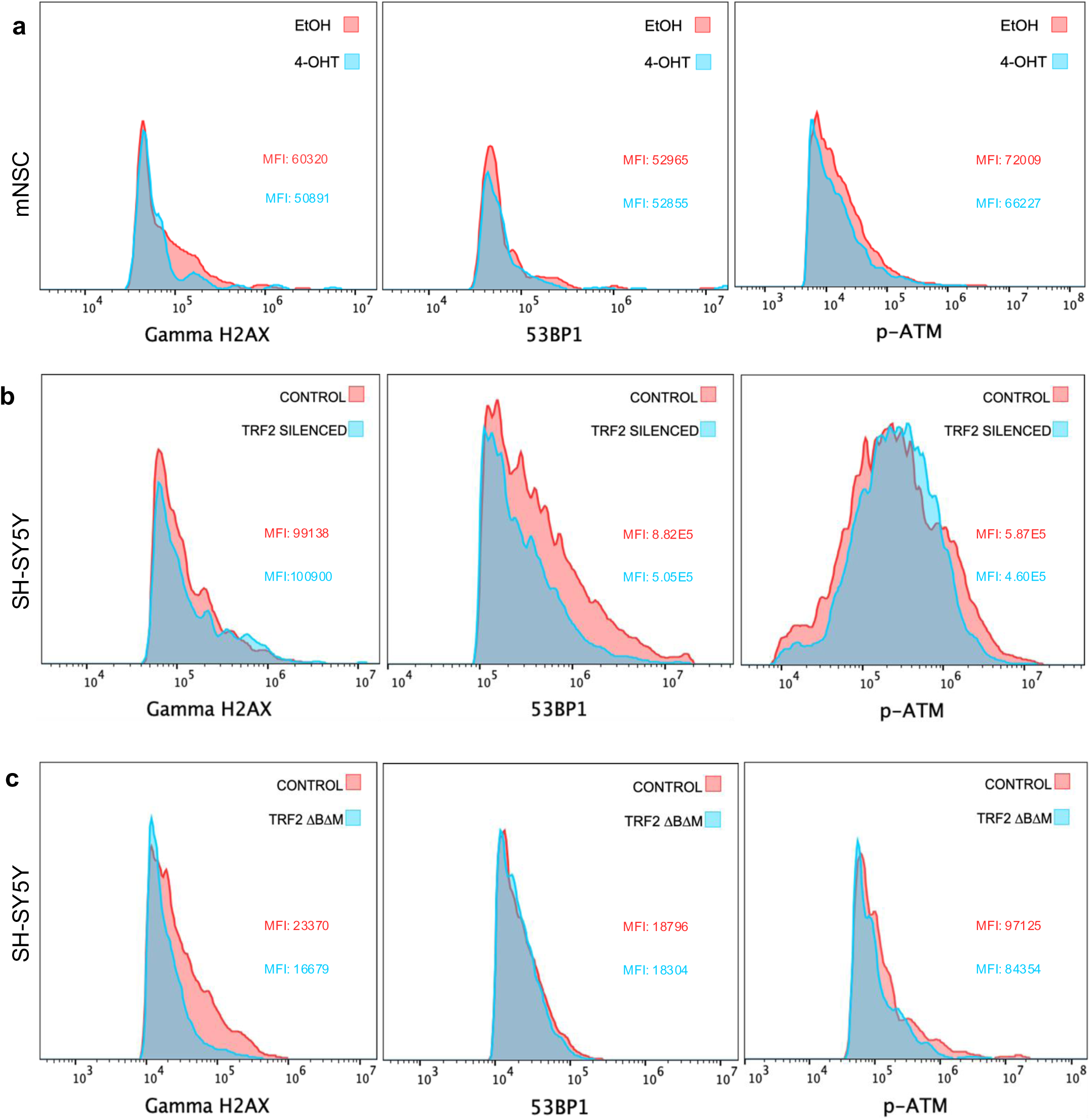
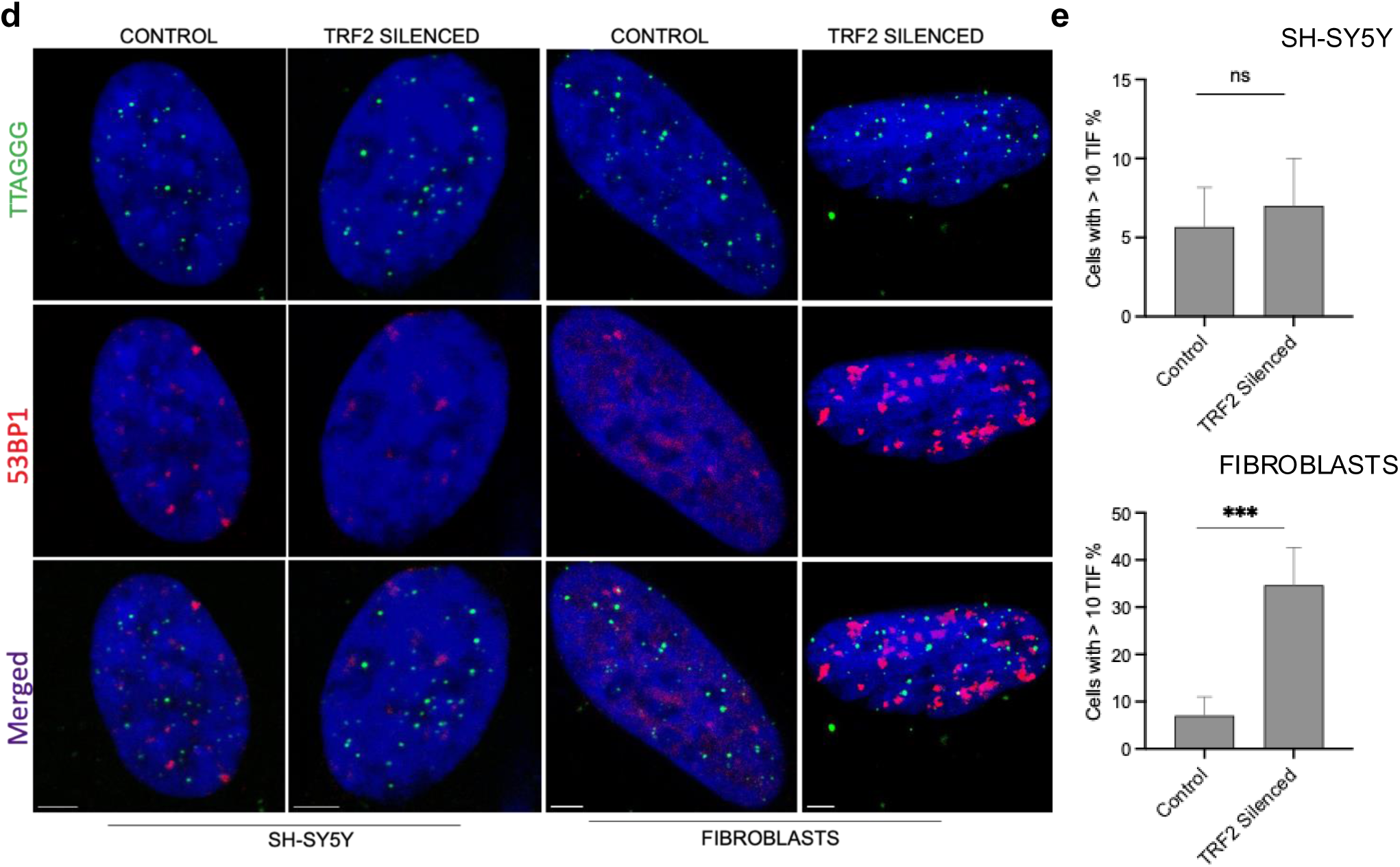
TRF2 mediated differentiation is independent of telomere damage. a. Flow cytometry with staining for γH2AX (i), 53BP1 (ii), p-ATM (iii) in mNSCs of *Terf2^F/F^*/*Nes:Cre* mice. Vehicle control (EtOH) and TRF2 knockout cells (4-OHT). Mean intensity of fluorescence is shown. The cell counts are normalized to respective modes for comparative representation b. Flow cytometry with staining for γH2AX (i), 53BP1 (ii), p-ATM (iii) in SH-SY5Y cells. Control and TRF2 silenced cells. Mean intensity of fluorescence (MIF) is shown. The cell counts are normalized to respective modes for comparative representation. c. Flow cytometry with staining for γH2AX (i), 53BP1 (ii), p-ATM (iii) in SH-SY5Y cells. Control and TRF2 ΔBΔM overexpression cells. Mean intensity of fluorescence (MIF) is shown. The cell counts are normalized to respective modes for comparative representation. d. Representative immunofluorescence and fluorescence in situ hybridization (IF–FISH) for 53BP1 (red) and telomeres (green) inSHSY5Y cells and Fibroblasts treated as indicated. Scale bar, (3 µm). e, Quantification of the percentage of cells with more than 10 53BP1 foci colocalizing with telomeres e. Quantification of the percentage of cells with more than 10 53BP1 foci colocalizing with telomeres, detected as in d. Error bars indicate standard deviation

To further test we induced double-strand breaks by treating SH-SY5Y with Doxorubicin (known to induce telomere dysfunction [37]). Enhanced telomere dysfunction-induced 53BP1 foci was clear at telomeres (Sup Fig 2b). To additionally ascertain telomere dysfunction induced DDR in SH-SY5Y cells, another shelterin component, Telomere Repeat binding Factor 1(TRF1), was silenced. This resulted in a sharp increase in □H2AX, in contrast to TRF2 depletion (Sup Fig 2c). Therefore, although telomere dysfunction induced DDR remains functional, this was not activated in response to TRF2 depletion. Together this show telomere protection function is conferred by TRF1, whereas TRF2 was dispensable for telomere protection in NSCs.

Next, we inspected whether differentiated neurons formed via loss of TRF2 were viable. For this, a dual-color fluorescent assay was used for the measurement of neurite outgrowth and cell viability (intracellular esterase activity [38]). Esterase activity assay and neurites indicated differentiated neurons were viable (Sup Fig 2d) substantiating TRF2 depletion does not induce cellular dysfunction, supporting absence of DDR. Taken together, these negate the possibility that telomere dysfunction induced DDR from TRF2 loss affects neural cell fate.

### TRF2 regulates transcription of key neurogenesis genes

TRF2 binding sites on potential G4-forming sequences from ChIP-seq (done earlier in HT1080 cells [18] showed enrichment of genes associated with neuronal differentiation (Sup Fig 3a). From these six genes with putative TRF2-G4 binding sites within promoters (Sup Fig 3b), and widely reported to promote neuronal differentiation were selected for further study: Amyloid precursor protein (APP) acts as a cell surface receptor facilitating neurite outgrowth [39–41]; Brain-derived neurotrophic factor (BDNF) promotes neurogenesis and neuronal differentiation by activation of signaling pathways that support neuronal growth and survival [42–44]; Glial cell line-derived neurotrophic factor family receptor alpha 1 (GFRA1) promotes presynaptic differentiation and synaptogenesis in hippocampal neurons [45,46]; L1 cell adhesion molecule (L1CAM) facilitates neurogenesis and neuronal differentiation by promoting neuronal migration, axon growth, and synaptic plasticity through interactions with other cell adhesion molecules [47–49]; Microtubule-associated protein 2 (MAP2) is crucial for neuronal differentiation by stabilizing microtubules and promoting dendritic morphogenesis [50,51]; and Presenilin-1 (PSEN1) was reported to promote neural differentiation [52,53] particularly by inducing the Netrin/DCC signaling pathway that enhances axon guidance and migration in cortical development [54] and cytoskeletal interactions [55].

**Figure 3:**
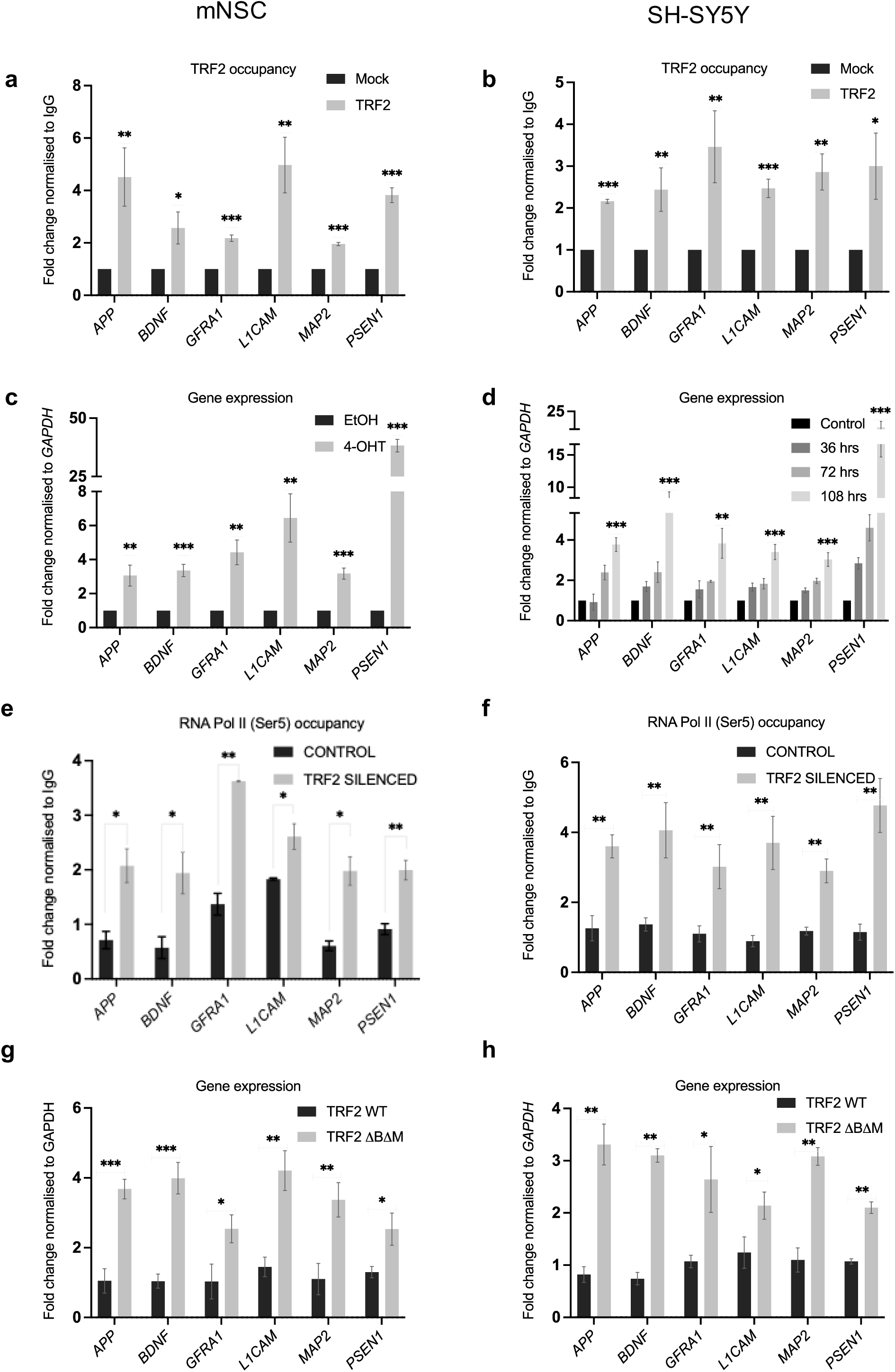
TRF2 regulates the transcription of neurogenesis genes. a. TRF2 ChIP followed by differentiation genes promoter qPCR for TRF2 binding in mNSCs (a) and SH-SY5Y (b) c. Gene expression changes upon tamoxifen (4-OHT) treated mNSC after 5 days. Fold change normalized over cells treated with vehicle control EtOH. d. Effect of TRF2 silencing on differentiation gene expression at increasing time points post transfection. Fold change normalized over cells treated with scrambled siRNA control. e. Pol2 (Ser5) occupancy on the promoters of differentiation genes following TRF2 silencing in (mNSCs (e) and SHSY5Y (f)). g. Gene expression changes upon DNA binding mutant expression of TRF2 in mNSC (g) and SH-SY5Y(h).

First, we aimed to determine whether TRF2 binds to the promoters of the six genes in neural cells. ChIP-PCR (primers spanning the G4 site in the near promoter as indicated for each gene in mouse/human (Sup Fig 3c)) confirmed promoter TRF2 binding in both mNSCs and human SH-SY5Y cells in all the six selected genes associated to neuronal differentiation (Fig 3a, b). Further upon TRF2 depletion, each of these six genes were upregulated in both mNSCs and SH-SY5Y supporting the role of TRF2 as a transcriptional repressor (Fig 3 c,d). In subsequent text we denote the selected six genes as TRF2-associated neuronal differentiation (TAN) genes.

Next, to validate the direct effect of TRF2 on regulation of the TAN genes, TRF2 was silenced, and RNA Pol-II (phospho-Ser5 initiation RNA polymerase) ChIP was performed. Loss of TRF2 significantly increased RNA Pol II occupancy on the promoters of genes associated to neuronal differentiation in both mNSCs and SH-SY5Y (Fig 3 e,f). Furthermore, induction of TRF2ΔBΔM, devoid of DNA binding, showed increase in expression of genes that had TRF2 occupancy in their promoters (Fig 3 g,h). Hence, both the basic and MYB domains of TRF2 are crucial for transcriptional regulation. Taken together, these results show direct DNA binding and transcriptional repression of the TAN genes (APP, BDNF, GFRA1, L1CAM, MAP2, PSEN1) by TRF2.

### Epigenetic histone modifications at promoters of neurogenesis genes are TRF2 mediated

TRF2 mediated histone methylations have been reported earlier [11]. H3K27me3 [56] and H3K9me2 [57] were noted to be predominant histone repressive marks for genes associated to neuronal differentiation. We sought to understand if TRF2 mediated repression influences the epigenetic state of promoters of the TAN genes. On TRF2 deletion, significant reduction in H3K27me3 occupancy within promoters of the TAN genes was clear in both mNSCs and SH-SY5Y (Fig. 4a, b). Whereas, the change in H3K9me2 occupancy was insignificant (Sup Fig 4a).

**Figure 4:**
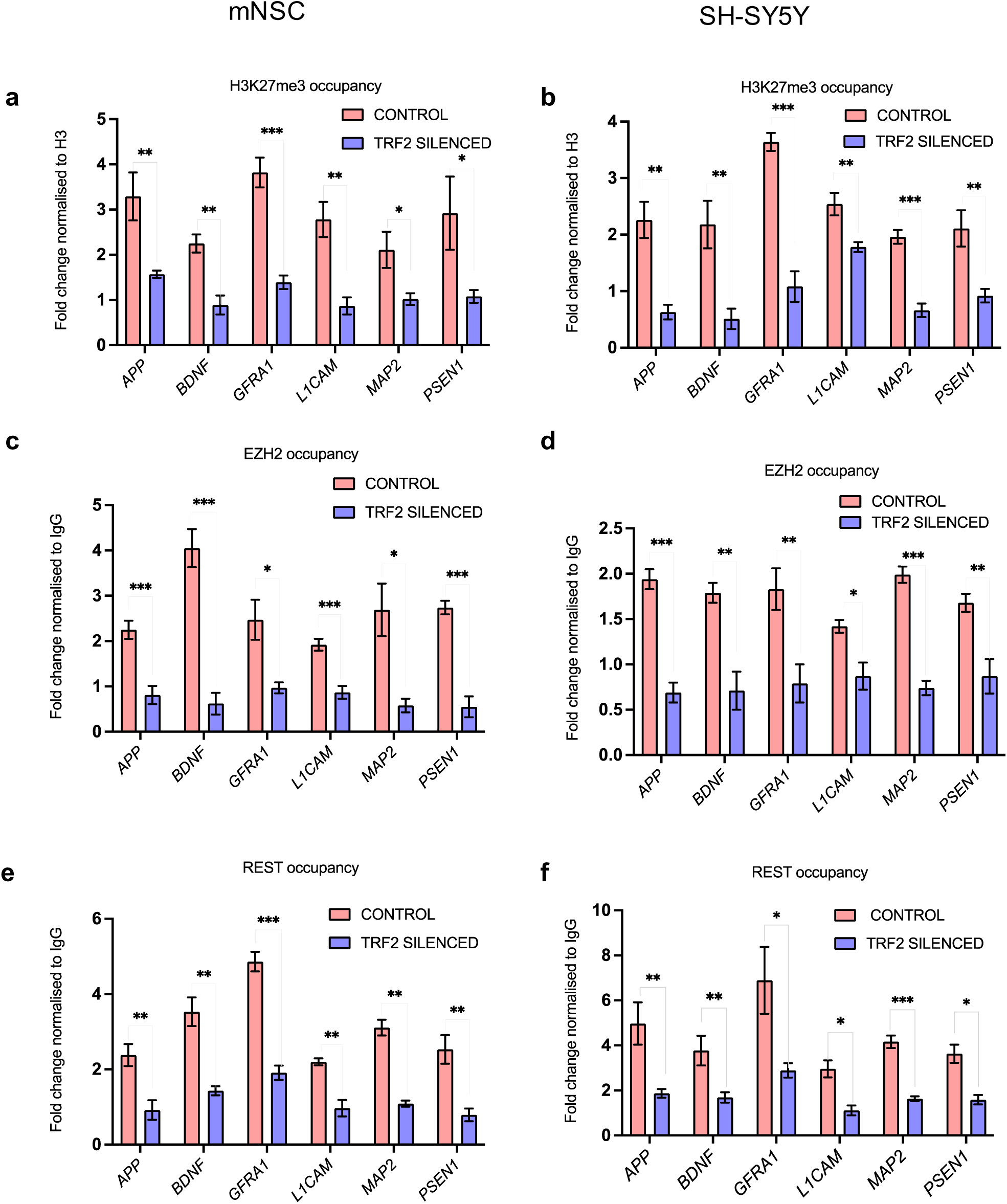
Epigenetic histone modifications at promoters of neurogenesis genes is TRF2 mediated. a. Effect of TRF2 silencing on H3K27me3 occupancy on the promoters of differentiation genes in mNSC (a) and SH-SY5Y (b). c. EZH2 occupancy on the promoters of differentiation genes on silencing TRF2 in mNSC (c) and SH-SY5Y (d). e. REST occupancy on the promoters of differentiation genes upon TRF2 silencing mNSC (e) and SH-SY5Y (f).

The Polycomb repressive complex 2 (PRC2), which includes the DNA methylation factor EZH2, catalyzes the trimethylation of H3K27 leading to gene suppression [58]. On TRF2 silencing, we found clear loss of EZH2 occupancy from promoters of the TAN genes in both mNSCs and SH-SY5Y (Fig 4 c,d).

Previous studies reported REST-dependent engagement of the PRC2 complex [59], and TRF2 was shown to interact with REST [60]. We asked whether TRF2 mediates REST recruitment to promoters of neuronal differentiation genes. On TRF2 silencing we observed reduction in REST occupancy at the TAN promoters in both mNSCs and SH-SY5Y (Fig 4 e,f). On the other hand, depletion of REST did not alter the occupancy of TRF2 on these promoters (Sup Fig 4b). Collectively, these show TRF2 recruits the REST-EZH2 complex, facilitating H3K27me3 deposition to repress the TAN genes.

### Post translational modification of TRF2 necessary for repressor function

Following this we sought to understand the role of TRF2 post translational modification(s) (PTM), if any, on neuronal differentiation. TRF2 PTM mutants (arginine methylation deficient R17H; acetylation deficient K176R, K190R or phosphorylation deficient T188N) were expressed in SHSY5Y (Sup Fig 5a). Of these, cells induced with the K176R mutant showed most significant increase in differentiation (Fig 5a). This was further ascertained with the increase in expression of the TAN genes (Fig 5b).

**Figure 5:**
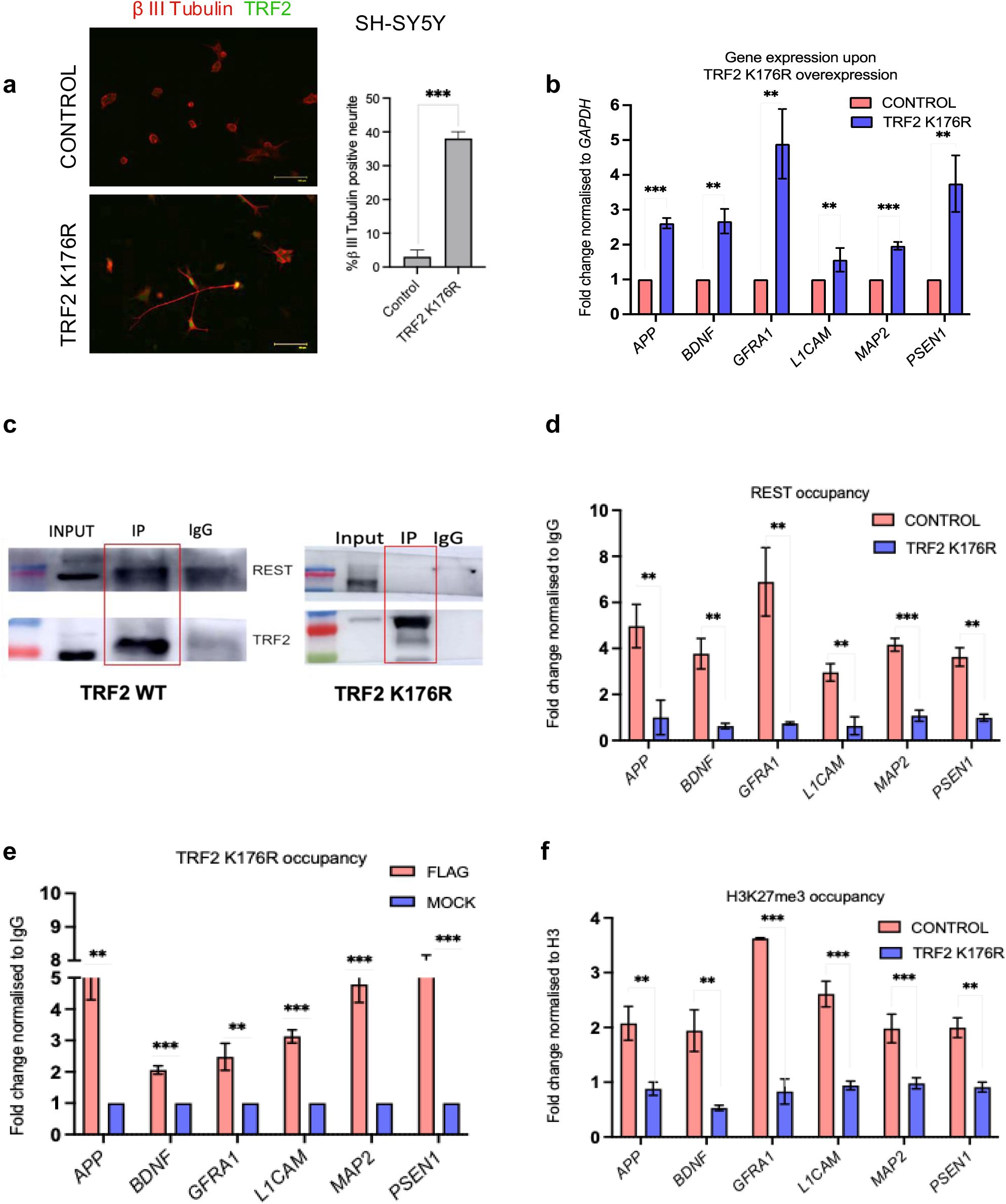
Post translational control of TRF2 affects its transcription factor activity. a. Immunofluorescence staining for TRF2 and β- III Tubulin proteins in Control and TRF2 K176R expression conditions. β- III Tubulin and TRF2 were stained using Alexa fluor-594 (red signal) and Alexa fluor-488 (green signal), respectively. Quantification of β- III Tubulin positive neurite from 30 cells (n = 30) shown in respective right panels (error bars ± SE). b. Effect of TRF2 K176R on differentiation gene expression 108 hrs post transfection. c. Co- Immunoprecipitation of WT TRF2 and K176R TRF2. d. REST occupancy on the promoters of differentiation genes in the presence of TRF2 K176R. e. Effect of TRF2 K176R on H3K27me3 occupancy on the promoters of differentiation genes. f. Occupancy of TRF2 K176R on the promoters of differentiation genes. ChIP for FLAG (TRF2).

TRF2’s dimerization and hinge domains (amino acids 45-446) were reported to be required for association with REST [60]. We therefore investigated whether the loss of function of TRF2 K176R was due to impaired interaction with REST. REST recruitment to the promoters of the TAN genes was reduced with the K176R mutation (Fig. 5d), along with decreased H3K27me3 enrichment at these promoter sites (Fig 5e).

The K176R mutation does not lie within DNA binding domains. Nevertheless, we sought to rule out the possibility that the loss of function was due to compromised DNA binding ability. In the FLAG-tagged TRF2 K176R overexpressed cells, ChIP with FLAG antibody was performed. The occupancy of the mutant TRF2 on the promoter indicated that the DNA binding function was intact (Fig 5f). Taken together, these results suggest that acetylation of TRF2 at K176 is crucial for interaction and recruitment of REST to promoters.

### TRF2 occupancy on the promoters of key neurogenesis genes is G-quadruplex dependent

We identified putative G4-forming sequences within promoters of the TAN genes in both human and mouse genomes (Sup fig 6a). First we checked if oligonucleotide sequences comprising the G4-forming sequences within the TAN promoters adopted the G4 structure in solution. Specific guanine base substitutions essential for G4 formation were introduced as negative controls (Sup Fig 6a). Both human and mouse G4 motifs displayed the characteristic parallel G4 form (noted using circular dichroism (CD) measurements), which was largely lost on the base substitutions (Sup Fig 6b).

**Figure 6:**
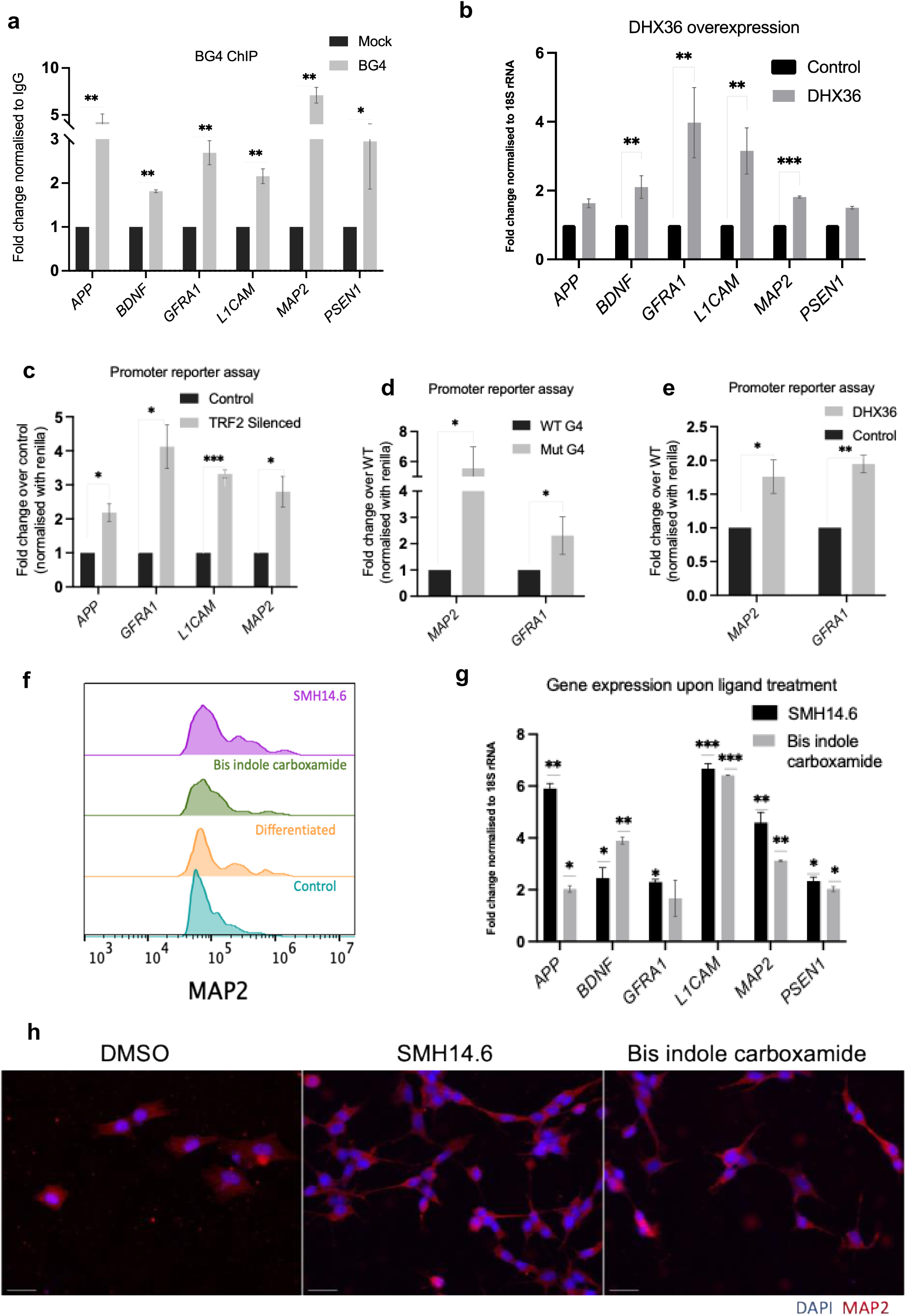
TRF2 occupancy on the promoters of neurogenesis genes is G-quadruplex dependent. a. BG4 ChIP followed by q-PCR for the promoters of genes associated to differentiation in SH-SY5Y cells. b. Expression of genes negatively regulated by TRF2 upon DHX36 overexpression. c. Quantification of luciferase expression driven by *APP, GFRA1, MAP2 and L1CAM* promoters upon TRF2 silencing d. Quantification of luciferase expression driven by *GFRA1, MAP2* promoters with the corresponding mutant G4. e. Quantification of luciferase expression driven by *GFRA1, MAP2* promoters with DHX36 expression. f. FACS based quantification of MAP2 promoter driven GFP expression upon treatment of ligands at 1.5 uM concentration for 5 days. g. Expression of genes regulated by TRF2 upon treatment of ligands at 1.5 uM concentration for 108hrs. h. Immunofluorescence staining for β- III Tubulin proteins in Control and Ligand treatment conditions for 7 days. β- III Tubulin was stained using Alexa fluor-594 (red signal). Scale bar 200 µm.

To see if the G4s were present inside cells we used ChIP with the reported G4-binding BG4 antibody [61]. In SH-SY5Y cells BG4 ChIP showed the occurrence of G4 in cells within promoters (ChIP-PCR with primers as shown in Sup Fig 3c) of each of the TAN genes (Fig 6a).

We further used the dead box helicase DHX36 reported for specific binding and resolution of promoter G4s [62]. In presence of DHX36 upregulation of all the TAN genes with promoter G4 motifs, and TRF2 occupancy, was evident underscoring the mechanistic significance TRF2-G4 binding (Fig 6b).

Further, luciferase promoter-reporter constructs were made with promoters of *MAP2, APP, L1CAM, and GFRA1*. On TRF2 silencing, all the luciferase reporters showed increase in promoter activity in SH-SY5Y cells confirming TRF2’s role as a transcriptional repressor (Fig 6c). In addition, to test the role of prompter G4s, G4-disrupting base substitutions were introduced within the *MAP2* and *GFRA1* promoter reporters. As expected, TRF2-dependent promoter repression was markedly impaired (Fig 6d). Furthermore, in presence of DHX36, luciferase reporter expression was enhanced in case of both *MAP2* and *GFRA*1 promoters (Fig 6e). Together, these confirm the role of intact G4s in TRF2-induced repression.

The functional relevance of the promoter G4s was further investigated by a cassette of GFP-reporter driven by *MAP2* promoter and integrated in SH-SY5Y cells (see methods). On TRF2 silencing enhanced luciferase activity from the integrated promoter cassette was clear. Interestingly, upon treatment using the G4 binding ligands SMH14.6 and Bis indole carboxamide in SH-SY5Y cells, GFP reporter activity increased (Fig 6f). Following this, we checked the effect of the ligands on TAN genes: All the TAN genes were upregulated following treatment with SMH14.6 and Bis indole carboxamide (Fig 6g). In addition, treatment with SMH14.6 and Bis indole carboxamide showed enhanced neurite outgrowth promoting neuronal differentiation (Fig 6h). These findings demonstrate the functional dependence of TRF2’s transcription repression of TAN genes to be dependent on G4s during neuronal differentiation.

## Discussion

Here, multiple lines of evidence uncover telomere-independent function of TRF2 in maintaining neural stemness. Non-telomeric TRF2: (a) directly binds to promoter DNA and b) recruits the REST-PRC2 epigenetic repressor complex to silence key neuronal differentiation genes. Recruitment of the REST-PRC2 complex by TRF2 was specific and dependent on TRF2 acetylation at the K176 residue. Interestingly, G4 motifs at the promoters of neuronal differentiation genes were required for TRF2 binding. Moreover, loss of TRF2 did not induce telomeric DNA damage or activate DDR as was evident from relatively unaltered levels of key DDR markers □H2AX, 53BP1, and p-ATM.

The lack of activation of DDR following TRF2 loss in NSCs aligns with findings showing TRF2 is not required for telomere protection in ESCs [3,4]. In ESCs, although depletion of TRF2 was reported to induce the two-cell embryo-like state, the role non-telomeric TRF2 was not studied [3]. Our findings from NSCs underline the importance of non-telomeric TRF2 as an epigenetic modulator of the stem cell state.

In proliferating neural stem cells, TRF2-mediated stabilization of REST was reported [60]. However, direct DNA-binding by TRF2 or TRF2-dependent recruitment of REST to promoters was not studied. TRF2-dependent recruitment of REST, and how this engages the canonical PRC2 repressor complex to deposit the H3K27me3 repressive mark to maintain neural stemness revealed here therefore underscores the critical and direct role of non-telomeric TRF2 in NSCs.

The direct involvement of DNA secondary structure G4s in DNA binding by TRF2 is interesting to note. Particularly, considering the subsequent epigenetic alterations through TRF2-mediated recruitment of epigenetic modifiers like REST and PRC2, which together underline the importance of non-duplex DNA in epigenetic control of the neural stem cell state. This was noted earlier although in a different context and cell type [15]. The displacement of TRF2 from gene promoters by G4-binding ligands, and the resulting induction of differentiation is notable. Although further work will be required to fully understand this, it underscores the possibility of ligand induced switching (on/off) of the neural stem state. On a similar note, function of the helicase DHX36 in promoting transcriptional activation (Fig 6b, 6e) suggests how specific helicases could be of interest in modulating the neurogenesis.

Being the first study on non-telomeric TRF2 and NSC, here we focused on specific neuronal differentiation markers to understand the role of TRF2 in transcription. This opens up the possibility of similar or related function at other potential TRF2 targets. Additionally, to first primarily understand molecular function we relied on in vitro models. While these help reveal the role of non-telomeric TRF2 in epigenetic control of neuronal differentiation, further work will be necessary to delineate how TRF2 affects NSC differentiation in vivo, and whether and how this might impact brain function. Additionally, RNA sequencing analysis (data not shown) indicated that glial markers were not significantly affected, suggesting a neurogenic shift potentially linked to compromised stemness maintenance. Future studies employing single-cell transcriptomics and lineage tracing could disentangle these effects by mapping TRF2’s transcriptional regulation to distinct NSC fates, providing deeper insights into its overall impact on neural stemness and differentiation in vivo.

Telomere shortening, a hallmark of aging, is notably evident in neurodegenerative diseases such as Alzheimer’s Disease (AD) [63,64]. Similar to other cell types, shortened telomeres affect the proliferative capacity of NSCs reducing the self-renewal potential of progenitor cells critical for normal adult neurogenesis [65–67]. Non-telomeric DNA binding and transcriptional function of TRF2 was noted to be telomere length dependent; furthermore, interestingly, this was found to affect different cellular contexts in a telomere-sensitive fashion [11,16,68,69]. RAP1, another shelterin factor, was also noted to bind non-telomeric sites across the genome [70–72]. Findings here, therefore, opens the possibility of non-telomeric TRF2 (and/or other shelterins) affecting neurogenesis in a way that is telomere sensitive. Together, providing molecular basis to neurodegeneration (and diseases like AD) on telomere shortening, or ageing.

In conclusion, this study identifies TRF2 as a critical regulator of neural stemness and neuronal differentiation through its non-canonical, telomere-independent transcriptional roles. By binding to G4 structures and recruiting REST and PRC2 repressive complexes, TRF2 maintains the neural stem state. These not only underscore the non-telomeric function of TRF2 in neurogenesis, but also open avenues to understand ageing associated neural pathophysiology in the light of new molecular mechanisms.

## Acknowledgements

We are thankful to all members of the S.C. laboratory for their suggestions/inputs and the IGIB Core Imaging Facility for its service facility. We acknowledge Dr. Hiyaa Ghosh and team for transgenic mouse generation and the NCBS animal facility for their support. We gratefully acknowledge the support of the Animal House Facility at the Institute of Genomics and Integrative Biology (IGIB) for providing the necessary resources and infrastructure, with experiments conducted under the approval of the Institutional Animal Ethics Committee (IAEC) vide approval number [IGIB/IAEC/10/Nov/2023/07]. Research fellowships to S.V., A.K.B.,S.B., (CSIR) are acknowledged. S.C. acknowledges support from Wellcome Trust/DBT India Alliance Fellowship (IA/S/18/2/504021) for research grants. Support from Council of Scientific and Industrial Research (CSIR) and Department of Biotechnology (DBT) to S.C. are also acknowledged.

## Author contributions

Conceptualization, S.C.; Methodology, S.V, A.K.B, S.B; Validation, S.V, A.K.B, S.B; Formal S.V.M., A.K.B, S.B, A.P, S.G, A.M, Analysis A.S.G.; Investigation, S.V, A.K.B, S.B.; Resources S.V, A.K.B, S.B.; Data Curation, S.V. A.K.B,; Writing – Original Draft Preparation, S.V; Writing – Review & Editing, S.C, S.V; Visualization, S.V.; Supervision, S.C.; Project Administration, S.V., S.C.; Funding Acquisition S.C.

## Declaration of interests

None

## Materials and methods

### Mice

All animals were housed, bred, and utilized in accordance with the protocols approved by the Institutional Animal Care and Ethics Committee. The *Terf2*F/F mice were obtained from the Jackson Laboratory (Strain #: 006568)[27]. The Nestin-CreERT2 transgenic line was generously provided by Dr. Hiyaa Ghosh from the National Centre for Biological Sciences, Bengaluru, India [28]. For conditional targeting of Terf2 in Nestin-expressing progenitors within the adult brain, the *Terf2*-flox line was crossed with the Nestin-CreERt2 line to generate Nestin-CreERt2; *Terf2*F/F (KO) or Nestin-CreERt2; *Terf2*+/+ (WT) mice. All animals were bred and housed at the Animal Care and Resource Centre.

### Cell line

SH-SY5Y and Fibrobalsts cells were cultured in Dulbecco’s Modified Eagle’s Medium-High Glucose (DMEM-HG) supplemented with 10% FBS. The differentiation of SH-SY5Y cells was conducted according to established methods [9]. Briefly, cells were plated at a concentration of 10^5 cells per mL on culture dishes coated with poly-D-lysine. They were then sequentially treated with 10 μM retinoic acid (RA; Sigma) in DMEM with serum for 5 days, followed by 10 ng/mL BDNF in serum-free medium for another 5 days. All cell lines were maintained at 37 degrees Celsius, with 5% CO2 and 95% humidity.

### Neurosphere culture

We adapted a previously published protocol for hippocampal neurosphere culture [28] as follows: Hippocampi from individual brains were micro dissected and digested in PDD [papain (2.5 U/ml), Dispase II (1 U/ml), and deoxyribonuclease I (10 μg/ml)] for 30 minutes in a shaker incubator at 37°C and 170 rpm, with intermittent trituration. The digestion was halted by adding complete neurobasal media (neurobasal supplemented with B27, GlutaMAX, and antibiotics).

The cell suspension was filtered through a 40-μm sieve and washed in wash buffer (30 mM glucose, 2 mM Hepes, and 26 mM NaHCO3 in 1× Hanks’ balanced salt solution) by centrifugation at 130g for 5 minutes at 21°C. The cell pellet was resuspended in 20 ml of complete neurobasal media, supplemented with EGF and FGF (20 ng/ml), and seeded into a flat-bottom 96-well plate. For Cre-induced deletion in neurosphere cultures, 0.5 μM 4-OH-Tmx (Sigma-Aldrich) was added to the cell suspension at the time of seeding. The cells were incubated in a tissue culture incubator with 5% CO2 for 5 to 10 days. Neurospheres were quantified at DIV7 and the results were expressed as the number of neurospheres per well for each brain.

### *MAP2* promoter driven GFP reporter cell line

The hMAP2 pGreenZeo Human Neurogenic Differentiation Reporter lentivirus (Cat.# SR10047PA-1) was procured from System Biosciences. Transduction of cells was performed following the manufacturer’s instructions and protocol. Post-transduction, cells were selected using zeocin at a concentration of 0.4 µg/ml. These selected cells were further serially passaged and maintained.

### ChIP (chromatin immunoprecipitation)

ChIP assays were conducted following the procedures outlined in Mukherjee et al. (2018)[11], utilizing specific primary antibodies Primary antibodies (TRF2 Novus Biologicals #NB110-57130, REST Sigma Aldrich #17-641, EZH2 Cell Signal Technology # 5246, H3K27me3 Abcam #ab6002, H3 Abcam # ab1791, H3K9me2 Abcam #ab1220 FLAG Merck #F3165) BG4 antibody Millipore Cat# MABE917; RNA PolII Ser5-P Abcam #ab5131) and IgG (from Millipore) as an isotype control. Three million cells were fixed with 1% formaldehyde, then lysed and chromatin was sheared to 200-300 bp fragments using a Biorupter (Diagenode). A 10% fraction of the sonicated chromatin was set aside as input, and processed using phenol-chloroform extraction and ethanol precipitation. ChIP was carried out using 3 mg of the appropriate antibody, and incubated overnight at 4°C. Immune complexes were captured using herring sperm DNA-saturated Magnetic Dynabeads (protein G/A) and extensively washed with low salt, high salt, and LiCl buffers. The Dynabeads were then resuspended in TE buffer and incubated with proteinase K at 55°C for 1 hour. DNA extraction was performed using phenol-chloroform-isoamyl alcohol, followed by precipitation with isopropanol, glycogen, and 3M sodium acetate. The resulting pellet was washed with 70% ethanol and resuspended in nuclease-free water. ChIP DNA validity was confirmed using q-PCR.

### Analysis of ChIP experiments

ChIP-qPCR analyses were conducted for TRF2, REST, and EZH2, using equal amounts of DNA quantified with the qubit HS DNA kit from each ChIP sample and its respective mock (IgG). Subsequently, the mean Ct values for each were utilized to determine fold change relative to IgG (mock). For histone ChIP assays, equal amounts of DNA from each histone ChIP and its respective total H3 ChIP were subjected to ChIP-qPCR. Fold change was calculated over total H3. In the case of BG4 ChIP, as the antibody was generated in E. coli, a mock control without antibody was utilized. ChIP-qPCR was performed with equal amounts of ChIP DNA and mock immunoprecipitation, and fold change over mock was determined similarly to other ChIP assays.

### Tel PCR

Tel PCR served as a positive control for TRF2 ChIP, given TRF2’s established role as a telomere-binding protein. Subsequently, following chromatin immunoprecipitation using an Anti-TRF2 antibody, Real-Time PCR was conducted with equal amounts of DNA template from both ChIP and IgG fractions. Telomere-specific primers (O’Callaghan et al., 2008)[73] were utilized, with an annealing temperature standardized at 52°C. The ChIP fraction readout was normalized against IgG to ascertain significant TRF2 binding at telomeres compared to background levels.

The primers used are:

Tel F CGGTTTGTTTGGGTTTGGGTTTGGGTTTGGGTTTGGGTT

Tel R GGCTTGCCTTACCCTTACCCTTACCCTTACCCTTACCCT

### Real time PCR for mRNA expression

RNA was isolated using TRIzol Reagent (Invitrogen, Life Technologies) following the manufacturer’s instructions. RNA quantification and cDNA synthesis were conducted using an Applied Biosciences kit. Quantitative real-time PCR was performed using a SYBR Green-based method (DSS TAKARA) to determine the relative transcript levels of mRNAs, with GAPDH serving as the housekeeping control gene for internal normalization. The average fold change was calculated by comparing the internally normalized threshold cycles (Ct) between the test and control samples. Each experiment was conducted in triplicate biological replicates, with technical triplicates of each primer pair and sample combination for q-PCR analysis.

### Co-Immunoprecipitation

Protein immunoprecipitation was conducted by harvesting six million cells (with overexpression either WT TRF2 or TRF2 K176R), then washed in cold 1X PBS and lysed using RIPA (Sigma) supplemented with 1x mammalian Protease Inhibitor Cocktail according to the manufacturer’s instructions. For the immunoprecipitation experiments, 1 mg of protein was incubated for 4 hours at 4°C with the primary antibody at the ratio recommended by the manufacturer for immunoprecipitation. The pull-down process was carried out using the catch-and-release co-immunoprecipitation kit (Millipore) following the manufacturer’s protocol.

### Immunofluorescence microscopy

Cells adherent to coverslips were cultured until reaching approximately 70% confluency. They were fixed using freshly prepared 4% Paraformaldehyde, and incubated for 10 minutes at room temperature (RT). Following fixation, cells were permeabilized with 0.2% Triton X-100 for 10 minutes at RT and then subjected to blocking with a solution containing 5% BSA in PBS for 2 hours at RT. This was followed by three washes with ice-cold PBS for 5 minutes each. After blocking, cells were treated with the appropriate antibodies: TRF2 Santa Cruz #sc52968, β-III Tubulin abcam # ab18207, PAX6 abcam #ab78545, 53BP1 abcam #ab36823 and incubated overnight at 4°C in a humid chamber. Following overnight incubation, cells were alternately washed with PBS and PBST three times and then probed with secondary antibodies (rabbit Alexa Fluor 488, 1:1000 / mouse Alexa Fluor 594, 1:1000) for 2 hours at RT. Cells were again washed alternately with PBS and PBST three times and mounted with Prolong Gold anti-fade reagent containing DAPI. Imaging was conducted using a EVOS M500 and Leica TCS-SP8 confocal microscope, signal intensities were calculated using ImageJsoftware post-ROI definitions.

Mouse brains at postnatal day P45 were collected after cardiac perfusion, and the brains were sectioned using a sledge microtome (Leica SMZ2010R) in frozen condition. Mouse brain sections were mounted on plus slides (Catalogue number: EMS 71869-11) and dried for 2– 3□hours. Slides were then transferred to a slide mailer (Catalogue number: EMS 71549-08) containing PBS□+□0.01% Triton X-100 for 10□minutes. This was followed by a 2× wash with PBS□+□0.03% Triton X-100 for 5□minutes. For antigen retrieval, sections were boiled in a 10□mM sodium citrate buffer (pH□=□6) at 90°C for 10□minutes using a water bath. Slides were then cooled to room temperature and washed with PBS□+□0.01% Triton X-100 for 10□minutes. Blocking was performed using 5% horse serum/Lamb serum in PBS□+□0.1% Triton X-100 for 1□hour, followed by overnight primary antibody incubation at 4□°C. Secondary antibody incubation was performed at room temperature for 2□hours. This was followed by three to five washes with 1× PBS. Slides were mounted using Vectashield mounting media (Vector Laboratories H-1000-10) and imaged using a Leica SP8 confocal microscope. Images in Supplementary XYZ represent a single representative plane of a Z-stacked image. Image processing was performed using ImageJ (FIJI) software.

### PNA Telomere Probe Hybridization

Lyophilized PNA probes (PNA BIO; F1009 TelC-FITC) were stored at -20°C until use. Before use, the tube was spun down, and 5 nmol of probe was resuspended in 100 µL formamide to prepare a 50 µM stock (∼250 µg/mL, 100×), heated at 55°C for 5 min, and stored in aliquots at - 70°C, protected from light. Thawed aliquots were reheated at 55°C for 5 min before use. The hybridization buffer contained 20 mM Tris (pH 7.4), 60% formamide, and 0.5% blocking reagent (Roche 11096176001) or 0.1 µg/mL salmon sperm DNA with 0.1% Tween-20. Wash solution contained 2× SSC with 0.1% Tween-20. Optional treatments included RNase A (100 µg/mL in PBS) and pepsin (0.005% in 10 mM HCl, prepared fresh and warmed to 37°C). Slides were prepared following standard fixation methods. FFPE sections underwent deparaffinization in xylene, followed by PBS washes (2×, 2 min each). Optional RNase treatment was performed at 37°C for 20 min, followed by PBS and water washes. If required, slides were treated with pepsin for 5 min at 37°C, followed by PBS washes. Dehydration was done using 70%, 85%, and 100% cold ethanol (2 min each), followed by air drying. For hybridization, 0.2 µL PNA probe was mixed with 20 µL hybridization buffer (500 nM final concentration). Both the slide and hybridization mix were preheated at 85°C for 5 min before the probe was applied. After incubation at 85°C for 10 min, slides were left at room temperature for 1 h in the dark with wet towels to prevent drying. Coverslips were removed by immersing slides in wash solution, followed by two washes at 55–60°C for 10 min each and a final wash at room temperature. DAPI staining was performed using a 1/750 dilution of 0.5 mg/mL DAPI in 2× SSC for 10 min, followed by sequential washes in 2× SSC, 1× SSC, and water (2 min each). Slides were dried by quick centrifugation, mounted with a coverslip, and imaged using a fluorescence microscope.

### Immuno-flow cytometry

One million cells per condition underwent fixation with 4% formaldehyde for 10 minutes at room temperature, followed by three washes with ice-cold PBS for 10 minutes each. Cells were permeabilized using pre-chilled 0.2% Triton for 10 minutes, followed by another three washes with ice-cold PBS for 5 minutes each. Primary antibodies (Ki67 abcam#ab15580, γH2AX CST #S139 (20E3), 53BP1 abcam #ab36823, p-ATM CST #S1981(D25E5), MAP2 abcam #ab32454, BDNF abcam #ab108319, L1CAM abcam #ab208155, and TRF2 Santa Cruz #sc52968/ Merck #4A794)were diluted in 1% BSA (in PBS) at a 1:250 volume ratio and incubated with the cells as a cocktail for 2 hours at room temperature. After three additional washes with ice-cold PBS (10 minutes each), secondary antibodies (rabbit Alexa Fluor 488 at 1:1000 dilution and mouse Alexa Fluor 594 at 1:1000 dilution in 1% BSA in PBS) were added to the cells and incubated at room temperature for 1 hour. Following three more washes with ice-cold PBS (10 minutes each), cells were resuspended in 0.5 mL of PBS and analyzed for fluorescence intensity using an Acuuri c6 flow cytometer in the FL1 (488 nm) and FL4 (647 nm) channels. The resulting FCS files were processed using FlowJo software (version 10).

### Transfections and TRF2 silencing

Cells were transfected in a 1:3 complex of FUGENE HD and DNA/RNA using protocols previously described for TRF2 WT and mutant mammalian expression plasmids. A TRF2 siRNA pool was used for TRF2 silencing at the 150 pMol concentration, keeping cells treated with the same concentration of scrambled RNA as the control, as described in Mukherjee et al. (2018, 2019a). Cells were incubated with the transfection complex for 12 hours in the media, after which a media change was given. Cells were given fresh media changes every 24 hours. To explore the differentiation phenotype, the plasmids were cultured in media supplemented with G418 for 5 days. For TRF2 silencing, transfection was performed every 36 hours to sustain the silencing effect.

### Western blotting

Protein extracts were prepared for Western blot analysis by suspending cell pellets in either RIPA buffer or passive lysis solution that had been treated with a 1x mammalian protease inhibitor cocktail. The proteins were then separated using 10% SDS-PAGE and transferred onto polyvinylidene difluoride membranes (Immobilon FL, Millipore). Following membrane blocking, primary antibodies including anti-TRF2 antibody (Novus Biological), anti-antibody (Abcam), anti-REST (Millipore), anti-EZH2 (CST), and anti-GAPDH antibody (Santa Cruz) were applied. Secondary antibodies, anti-mouse, and anti-rabbit HRP conjugates, were sourced from CST. Finally, the blot was developed using the Millipore HRP chemiluminescence detection kit, and images were captured using a GE chemiluminescence imager.

### Luciferase assay

Promoters of genes associated with differentiation (up to 1500 bp upstream of the transcription start site) were individually cloned into the PGL3 basic vector upstream of a firefly luciferase construct. G4 motif-disrupting mutations were introduced into the promoter constructs via site-directed mutagenesis. A plasmid (pGL4.73) containing a CMV promoter driving Renilla luciferase was co-transfected as a transfection control for normalization. After 48 hours, cells were harvested, and the luciferase activities of the cell lysate were measured using a dual-luciferase reporter assay system (Promega).

### Circular dichroism

CD profiles (220–300 nm) were acquired for representative G4 motifs located within 50 bp of the Transcription Start Site (TSS) of a gene, which overlapped with TRF2 high-confidence ChIP seq peaks. A depiction of oligonucleotides and their corresponding mutated sequences is provided in Fig. 3A. CD analysis revealed the presence of a G4 motif with the unmodified sequence, while a mutated G4 sequence exhibited partial or complete disruption of the G4 motif under similar conditions (buffer composition for G-quadruplex formation: 10 mm sodium cacodylate and 100 mm KCl). The CD spectra were recorded using a Jasco-89 spectropolarimeter equipped with a Peltier temperature controller. The experiments were conducted in a 1-mm path-length cuvette across a wavelength range of 200–320 nm. Oligonucleotides were commercially synthesized by Sigma-Aldrich. Oligonucleotides at a concentration of 2.5 μm were diluted in sodium cacodylate buffer (10 mm sodium cacodylate and 100 mm KCl, pH 7.4) and denatured by heating to 95 °C for 5 min, followed by gradual cooling to 15 °C over several hours. The CD spectra presented herein represent the average of three scans acquired at 20 °C and have been adjusted to correct for signal contributions from the buffer.

### Sample processing, generation of cell-suspension and single-cell RNA sequencing (scRNA-Seq)

The freshly harvested mouse brain hippocampal tissue was homogenized using 1 ml of the brain dissociation medium (Papain [20 U/ml] + DNase I solution [100 µl/ml] + 0.5% BSA + modified HBSS) and incubated at 37°C for 30 minutes. The dissociated tissue was passed through 70 µm strainer and sample preparation medium (2% FBS + 1mM EDTA + 5% BSA + modified HBSS) was added. The solution was centrifuged at 400 rcf for 10 minutes followed by washing using 1 ml ice-cold DPBS. The solution was mixed with 500 µl freshly prepared isotonic percoll solution and 2 ml DPBS followed by centrifugation at 3000 rcf for 10 minutes which resulted in upper DPBS and lower percoll layer. The two layers were separated by myelin and debris, while cells were at the bottom. The layers were aspirated out leaving 500 µl having the cells. 4 ml DPBS was added and the cells were then centrifuged at 400 rcf for 10 minutes at 4°C followed by resuspension in the Hibernate media. The cells were counted using Hemocytometer and proceeded for single-cell RNA sequencing as prescribed by the manufacturer (CG000315 Chromium Next GEM Single Cell 3’ Gene Expression v3.1 Dual Index RevF, 10X Genomics). The sequencing of the library was done in NovaSeq 6000 (Illumina) using paired-end sequencing chemistry and run recipe of 28:10:10:90 (R1:I1:I2:R2).

### Analysis of single-cell RNA sequencing (scRNA-Seq) dataset

The raw sequence intensity files (BCL) were demultiplexed and converted into FASTQ files using the CellRanger v7.1.0 (https://www.10xgenomics.com/support/software/cell-ranger/latest) software suite (10X Genomics) mkfastq pipeline[74]. The quality of demultiplexed FASTQs were checked using FASTQC (https://github.com/s-andrews/FastQC) software [75]. The demultiplexed, quality checked FASTQs were then aligned to the mouse reference genome (https://cf.10xgenomics.com/supp/cell-exp/refdata-gex-GRCm39-2024-A.tar.gz) using the CellRanger v7.1.0 count pipeline[74]. The cell × gene count matrix was further processed using the R package Seurat v4.4.0 (https://github.com/satijalab/seurat)[76]. After removing cells expressing less than 200 genes, cells expressing more than 8000 genes and cells with mitochondrial reads more than 20%; good quality cells were clustered using Louvain algorithm and visualized using Uniform Manifold Approximation Projection (UMAP) using resolution 1.0. Doublets were removed from the pre-processed data using the R package DoubletFinder v2.0.3 (https://github.com/chris-mcginnis-ucsf/DoubletFinder)[77]. The marker genes for each cell cluster were identified using Seurat FindAllMarkers() which essentially utilizes the Wilcoxon rank-sum test and clusters were annotated to each cell type accordingly3. From the overall cell types identified, neurons were subclustered and expression of genes of interest were determined using Seurat DotPlot() and FeaturePlot()[76].

### Analysis of spatial and single-cell open-source transcriptomics data

Adult hippocampal single nuclear RNA seq data set from Habib et al., 2016 [23](GSE84371) was analyzed and graphs were generated using the single cell portal [78]. Original classification of clusters was retained as per Habib et al., 2016 [23]. Spatial transcriptomics data set was analyzed using Loupe browser V8.1.2. Dentate gyrus was defined by the expression of Prox1 and CA field was demarcated by expression of Dkk3. Log normalized expression value of Terf2 was plotted. (https://www.10xgenomics.com/support/software/loupe-browser). Data sets from Di Bella et al., 2021 (GSE153164) [24] was analyzed using single cell portal. Wheel plots for Terf2 was generated from La mano et al., 2021 (PRJNA637987) [25] using a webtool (http://mousebrain.org/wheel). 1.3 million brain cells from E18 mouse brain data from 10x genomics was analyzed using Loupe browser V8.1.2.

### Quantification and statistical analysis

Each experimental procedure was conducted in a minimum of three biological replicates. Means and standard errors were calculated in Excel. All statistical analyses and graphical representations were generated using GraphPad Prism V8.0.2. To compare two conditions, such as control versus TRF2 silenced, paired or unpaired Student’s T-tests were employed (*p < 0.05, **p < 0.01, ***p < 0.005, ****p < 0.0001).

**Table 1:**
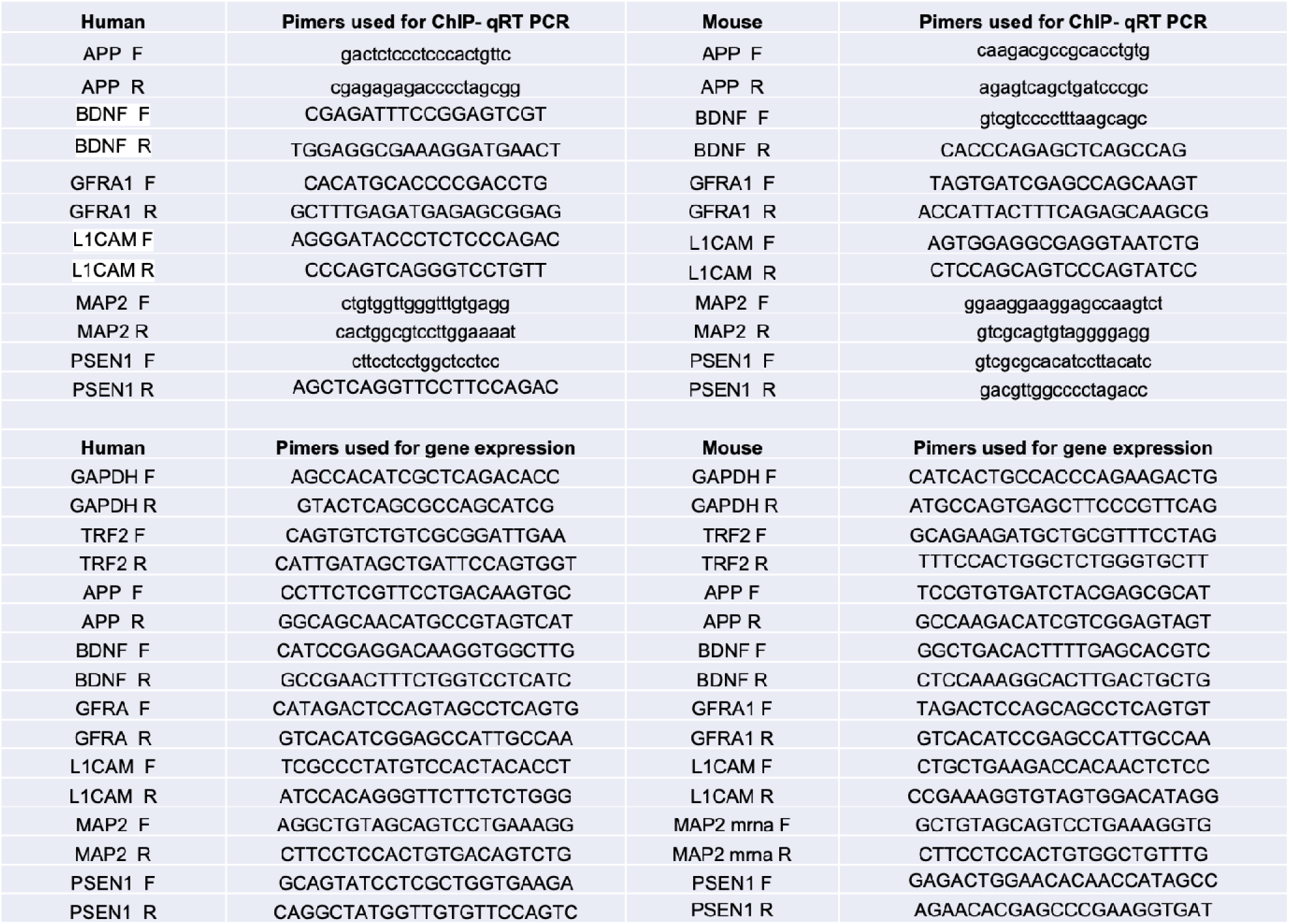
List of primers used.

## Notes

### Competing Interest Statement

The authors have declared no competing interest.

## References

1. Karlseder, J. et al. (1999) p53- and ATM-dependent apoptosis induced by telomeres lacking TRF2. Science (1979) 283, 1321–1325

2. Karlseder, J. et al. (2004) The Telomeric Protein TRF2 Binds the ATM Kinase and Can Inhibit the ATM-Dependent DNA Damage Response. PLoS Biol 2, e240

3. Markiewicz-Potoczny, M. et al. (2021) TRF2-mediated telomere protection is dispensable in pluripotent stem cells. Nature 589, 110–115

4. Ruis, P. et al. (2021) TRF2-independent chromosome end protection during pluripotency. Nature 589, 103–109

5. Wagner, K.D. et al. (2017) The differential spatiotemporal expression pattern of shelterin genes throughout lifespan. Aging 9, 1219–1232

6. Ovando-Roche, P. et al. (2014) TRF2-Mediated Stabilization of hREST4 Is Critical for the Differentiation and Maintenance of Neural Progenitors. Stem Cells 32, 2111–2122

7. Ying, Y. et al. (2022) The non-telomeric evolutionary trajectory of TRF2 in zebrafish reveals its specific roles in neurodevelopment and aging. Nucleic Acids Res 50, 2081–2095

8. Lobanova, A. et al. (2017) Different requirements of functional telomeres in neural stem cells and terminally differentiated neurons. Genes Dev 31, 639– 647

9. Zhang, P. et al. (2011) Nontelomeric splice variant of telomere repeat-binding factor 2 maintains neuronal traits by sequestering repressor element 1-silencing transcription factor. Proc Natl Acad Sci U S A 108, 16434–9

10. El Maï, M., et al. (2014) The telomeric protein TRF2 regulates angiogenesis by binding and activating the PDGFR$β$ promoter. Cell Rep 9, 1047–1060

11. Mukherjee, A.K. et al. (2018) Telomere length-dependent transcription and epigenetic modifications in promoters remote from telomere ends. PLoS Genet 14

12. Zizza, P. et al. (2019) TRF2 positively regulates SULF2 expression increasing VEGF-A release and activity in tumor microenvironment. Nucleic Acids Res 47, 3365–3382

13. Cherfils-Vicini, J. et al. (2019) Cancer cells induce immune escape via glycocalyx changes controlled by the telomeric protein TRF2. EMBO J 38, e100012

14. Hussain, T. et al. (2017) Transcription regulation of CDKN1A (p21/CIP1/WAF1) by TRF2 is epigenetically controlled through the REST repressor complex. Sci Rep 7, 1–13

15. S, S., et al. (2021) Human telomerase is directly regulated by non-telomeric TRF2-G-quadruplex interaction. Cell Rep 35

16. Mukherjee, A.K. et al. (2024) Telomere length sensitive regulation of Interleukin Receptor 1 type 1 (IL1R1) by the shelterin protein TRF2 modulates immune signalling in the tumour microenvironment. Elife 13

17. Purohit, G. et al. (2018) Extratelomeric Binding of the Telomere Binding Protein TRF2 at the PCGF3 Promoter Is G-Quadruplex Motif-Dependent. Biochemistry 57, 2317–2324

18. Mukherjee, A.K. et al. (2019) Telomere repeat-binding factor 2 binds extensively to extra-telomeric G-quadruplexes and regulates the epigenetic status of several gene promoters Journal of Biological Chemistry, 294 American Society for Biochemistry and Molecular Biology Inc., 17709–17722

19. Huppert, J.L. and Balasubramanian, S. (2007) G-quadruplexes in promoters throughout the human genome. Nucleic Acids Res 35, 406

20. Verma, A. et al. (2008) Genome-wide computational and expression analyses reveal G-quadruplex DNA motifs as conserved cis-regulatory elements in human and related species. J Med Chem 51, 5641–5649

21. Halder, R. et al. (2010) Guanine quadruplex DNA structure restricts methylation of CpG dinucleotides genome-wide. Mol Biosyst 6, 2439–2447

22. Mukherjee, A.K. et al. (2019) Non-duplex G-Quadruplex Structures Emerge as Mediators of Epigenetic Modifications. Trends Genet 35, 129

23. Habib, N. et al. (2016) Div-Seq: Single-nucleus RNA-Seq reveals dynamics of rare adult newborn neurons. Science (1979) 353, 925–928

24. Di Bella, D.J., et al. (2021) Molecular logic of cellular diversification in the mouse cerebral cortex. Nature 2021 595:7868 595, 554–559

25. La Manno, G., et al. (2021) Molecular architecture of the developing mouse brain. Nature 596, 92–96

26. Grammatikakis, I. et al. (2016) The long and the short of TRF2 in neurogenesis. Cell Cycle 15, 3026

27. Celli, G.B. and de Lange, T. (2005) DNA processing is not required for ATM-mediated telomere damage response after TRF2 deletion. Nat Cell Biol 7, 712–718

28. Shariq, M. et al. (2021) Adult neural stem cells have latent inflammatory potential that is kept suppressed by Tcf4 to facilitate adult neurogenesis. Sci Adv 7

29. Encinas, M. et al. (2000) Sequential treatment of SH-SY5Y cells with retinoic acid and brain-derived neurotrophic factor gives rise to fully differentiated, neurotrophic factor-dependent, human neuron-like cells. J Neurochem 75, 991–1003

30. Kovalevich, J. and Langford, D. (2013) Considerations for the Use of SH-SY5Y Neuroblastoma Cells in Neurobiology. Methods Mol Biol 1078, 9

31. Teppola, H. et al. (2015) Morphological Differentiation Towards Neuronal Phenotype of SH-SY5Y Neuroblastoma Cells by Estradiol, Retinoic Acid and Cholesterol. Neurochem Res 41, 731

32. Shipley, M.M. et al. (2016) Differentiation of the SH-SY5Y Human Neuroblastoma Cell Line. J Vis Exp 2016, 53193

33. Van Steensel, B., et al. (1998) TRF2 protects human telomeres from end-to-end fusions. Cell 92, 401–413

34. Hanaoka, S. et al. (2005) Comparison between TRF2 and TRF1 of their telomeric DNA-bound structures and DNA-binding activities. Protein Science 14, 119–130

35. Fouché, N. et al. (2006) The Basic Domain of TRF2 Directs Binding to DNA Junctions Irrespective of the Presence of TTAGGG Repeats. Journal of Biological Chemistry 281, 37486–37495

36. Mender, I. and Shay, J. (2015) Telomere Dysfunction Induced Foci (TIF) Analysis. Bio Protoc 5

37. Li, P. et al. (2012) Telomere dysfunction induced by chemotherapeutic agents and radiation in normal human cells. Int J Biochem Cell Biol 44, 1531–1540

38. Hancock, M.K. et al. (2015) A Facile Method for Simultaneously Measuring Neuronal Cell Viability and Neurite Outgrowth. Curr Chem Genom Transl Med 9, 6

39. Nicolas, M. and Hassan, B.A. (2014) Amyloid precursor protein and neural development. Development 141, 2543–2548

40. Baumkötter, F. et al. (2014) Amyloid precursor protein dimerization and synaptogenic function depend on copper binding to the growth factor-like domain. J Neurosci 34, 11159–11172

41. Wang, S. et al. (2016) Amyloid β precursor protein regulates neuron survival and maturation in the adult mouse brain. Molecular and Cellular Neuroscience 77, 21–33

42. Ahmed, S. et al. (1995) BDNF enhances the differentiation but not the survival of CNS stem cell-derived neuronal precursors. J Neurosci 15, 5765–5778

43. Rossi, C. et al. (2006) Brain-derived neurotrophic factor (BDNF) is required for the enhancement of hippocampal neurogenesis following environmental enrichment. European Journal of Neuroscience 24, 1850–1856

44. Gonzalez, A., et al. (2016) Cellular and molecular mechanisms regulating neuronal growth by brain-derived neurotrophic factor. Cytoskeleton 73, 612–628

45. Pozas, E. and Ibáñez, C.F. (2005) GDNF and GFRalpha1 promote differentiation and tangential migration of cortical GABAergic neurons. Neuron 45, 701–713

46. Ledda, F. et al. (2007) GDNF and GFRalpha1 promote formation of neuronal synapses by ligand-induced cell adhesion. Nat Neurosci 10, 293–300

47. Kiryushko, D. et al. (2004) Regulators of neurite outgrowth: role of cell adhesion molecules. Ann N Y Acad Sci 1014, 140–154

48. Schäfer, M.K.E. and Frotscher, M. (2012) Role of L1CAM for axon sprouting and branching. Cell Tissue Res 349, 39–48

49. Sytnyk, V. et al. (2017) Neural Cell Adhesion Molecules of the Immunoglobulin Superfamily Regulate Synapse Formation, Maintenance, and Function. Trends Neurosci 40, 295–308

50. Przyborski, S.A. and Cambray-Deakin, M.A. (1995) Developmental regulation of MAP2 variants during neuronal differentiation in vitro. Developmental Brain Research 89, 187–201

51. Harada, A. et al. (2002) MAP2 is required for dendrite elongation, PKA anchoring in dendrites, and proper PKA signal transduction. J Cell Biol 158, 541

52. Capell, A. et al. (1997) Cellular expression and proteolytic processing of presenilin proteins is developmentally regulated during neuronal differentiation. J Neurochem 69, 2432–2440

53. Flood, F. et al. (2004) Presenilin expression during induced differentiation of the human neuroblastoma SH-SY5Y cell line. Neurochem Int 44, 487–496

54. Wines-Samuelson, M. et al. (2005) Role of presenilin-1 in cortical lamination and survival of Cajal-Retzius neurons. Dev Biol 277, 332–346

55. Pigino, G. et al. (2001) Presenilin-1 Mutations Reduce Cytoskeletal Association, Deregulate Neurite Growth, and Potentiate Neuronal Dystrophy and Tau Phosphorylation. The Journal of Neuroscience 21, 834

56. Buontempo, S. et al. (2022) EZH2-Mediated H3K27me3 Targets Transcriptional Circuits of Neuronal Differentiation. Front Neurosci 16, 814144

57. Fiszbein, A. and Kornblihtt, A.R. (2016) Histone methylation, alternative splicing and neuronal differentiation. Neurogenesis 3

58. Margueron, R. and Reinberg, D. (2011) The Polycomb complex PRC2 and its mark in life. Nature 2011 469:7330 469, 343–349

59. Dietrich, N. et al. (2012) REST–Mediated Recruitment of Polycomb Repressor Complexes in Mammalian Cells. PLoS Genet 8, e1002494

60. Zhang, P. et al. (2008) Nontelomeric TRF2-REST Interaction Modulates Neuronal Gene Silencing and Fate of Tumor and Stem Cells. Current Biology 18, 1489–1494

61. Biffi, G. et al. (2013) Quantitative Visualization of DNA G-quadruplex Structures in Human Cells. Nat Chem 5, 182

62. Marshall, P.R. et al. (2024) DNA G-Quadruplex Is a Transcriptional Control Device That Regulates Memory. The Journal of Neuroscience 44, e0093232024

63. Blasco, M.A. (2007) Telomere length, stem cells and aging. Nat Chem Biol 3, 640–649

64. Hou, Y. et al. (2019) Ageing as a risk factor for neurodegenerative disease. Nat Rev Neurol 15, 565–581

65. Ferrón, S.R. et al. (2009) Telomere shortening in neural stem cells disrupts neuronal differentiation and neuritogenesis. J Neurosci 29, 14394–14407

66. Palmos, A.B. et al. (2020) Telomere length and human hippocampal neurogenesis. Neuropsychopharmacology 45, 2239

67. Harley, J. et al. (2024) Telomere shortening induces aging-associated phenotypes in hiPSC-derived neurons and astrocytes. Biogerontology 25, 341–360

68. Vinayagamurthy, S. et al. (2023) Telomeres expand sphere of influence: emerging molecular impact of telomeres in non-telomeric functions. Trends in Genetics 39, 59–73

69. Sengupta, A., et al. (2024) Telomeres control human telomerase (hTERT) expression through non-telomeric TRF2. bioRxiv DOI: 10.1101/2023.10.09.561466

70. Teo, H. et al. (2010) Telomere-independent Rap1 is an IKK adaptor and regulates NF-κB-dependent gene expression. Nat Cell Biol 12, 758–767

71. Crabbe, L. and Karlseder, J. (2010) Mammalian Rap1 widens its impact Nature Cell Biology, 12 Nature Publishing Group, 733–735

72. Zhang, X. et al. (2019) Telomere-dependent and telomere-independent roles of RAP1 in regulating human stem cell homeostasis. Protein Cell 10, 649–667

73. O’Callaghan, N.J. et al. (2008) A quantitative real-time PCR method for absolute telomere length. Biotechniques 44, 807–809

74. Zheng, G.X.Y. et al. (2017) Massively parallel digital transcriptional profiling of single cells. Nature Communications 2017 8:1 8, 1–12

75. Andrews S. (2010) FastQC: a quality control tool for high throughput sequence data. Available online at: http://www.bioinformatics.babraham.ac.uk/projects/fastqc

76. Hao, Y. et al. (2020) Integrated analysis of multimodal single-cell data. DOI: 10.1101/2020.10.12.335331

77. McGinnis, C.S. et al. (2019) DoubletFinder: Doublet Detection in Single-Cell RNA Sequencing Data Using Artificial Nearest Neighbors. Cell Syst 8, 329–337.e4

78. Tarhan, L. et al. Single Cell Portal: an interactive home for single-cell genomics data. DOI: 10.1101/2023.07.13.548886

