## Supplementary Figures for "TRF2 Non-Telomeric Function is Indispensable for Neural stemness"

**Supplementary Figure S1.1**

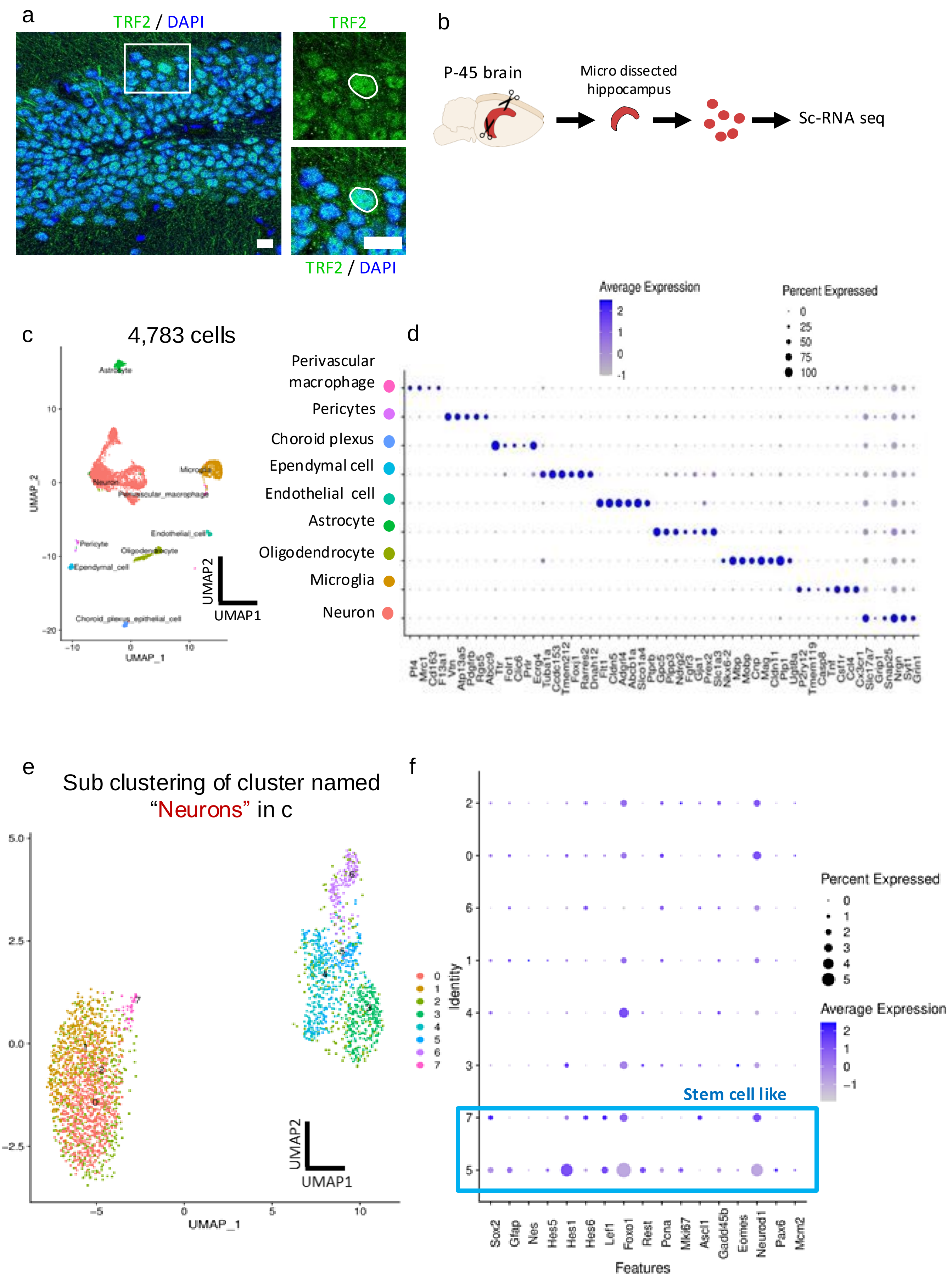

g

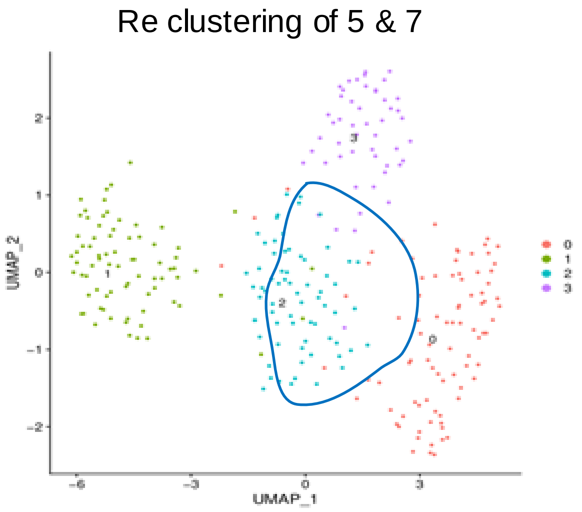


h

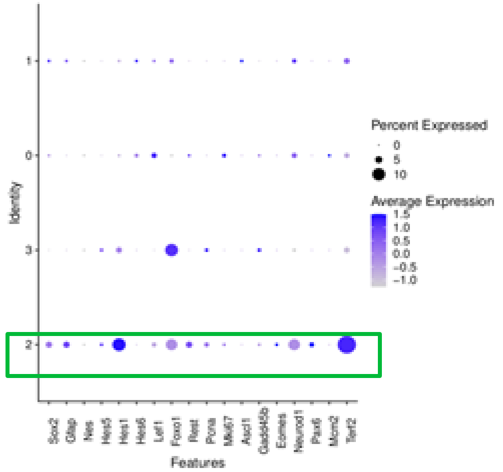


a. Representative confocal image showing nuclear localization of TRF2 in the Dentate Gyrus (DG) of the Hippocampus of a P45 wild type mouse brain. (Scale bars: 10μm)

b. Schematic of the sample processing for sc-RNA seq.

c. Annotated UMAP plot showing the nine major clusters from 4,784 cells obtained from the p45 wt mouse hippocampus.

d. Bubble plot displaying the marker gene expression for each defined cluster.

e. Subclusters obtained after re-clustering the cluster labeled "neuron" in (b).

f. Bubble plot showing the expression of marker genes for adult hippocampal neural stem cells (nscs), indicating that subclusters 5 and 7 exhibit stem cell-like signatures (blue box).

g. UMAP plot showing the re-clustering of subclusters 5 and 7.

h. Expression profile of *terf2* in the subclusters, showing high expression in cluster 2 (red box), which also exhibits significant expression of several stem cell markers (green box).

**Supplementary Figure S1.2**

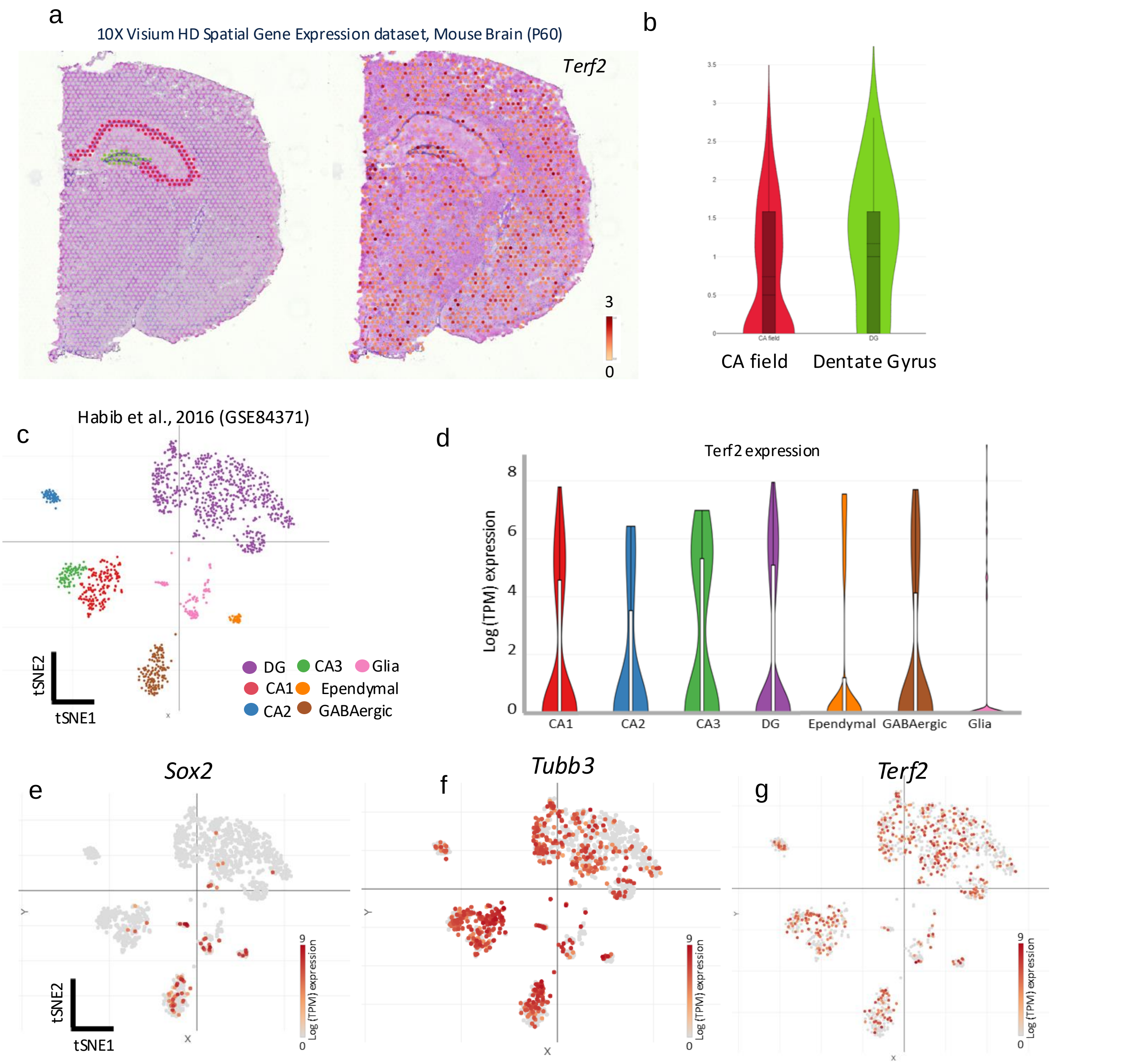


a. Publically available spatial transcriptomic data set reveals Terf2 transcripts are present in both Dentate gyrus and CA field of the hippocampus (Data set link: www.10xgenomics.com/datasets/visium-hd-cytassist-gene-expression-mouse-brain-fresh-frozen)

b. Violin plot showing log normalized expression levels of *Terf2* in the regions designated as DG and CA field. (D-H) Single Nucleus RNA seq (Habib et al., 2016, GEO accession no. GSE84371) of adult mouse hippocampus shows ubiquitous presence of Terf2 in the entire hippocampal cell population including DG.

c. tSNE plot & (d) Violin plot showing the expression profile of *Terf2* in different hippocampal subclusters.

e-g. Feature plots showing *Terf2* is present in both Tubb3+ and Sox2 + cells.
(Data set link: https://singlecell.broadinstitute.org/single_cell/study/SCP1/-single-nucleus-rna-seq-of-cell-diversity-in-the-adult-mouse-hippocampus-snuc-seq#study-visualize).

**Supplementary Figure Figure S1.3**

a

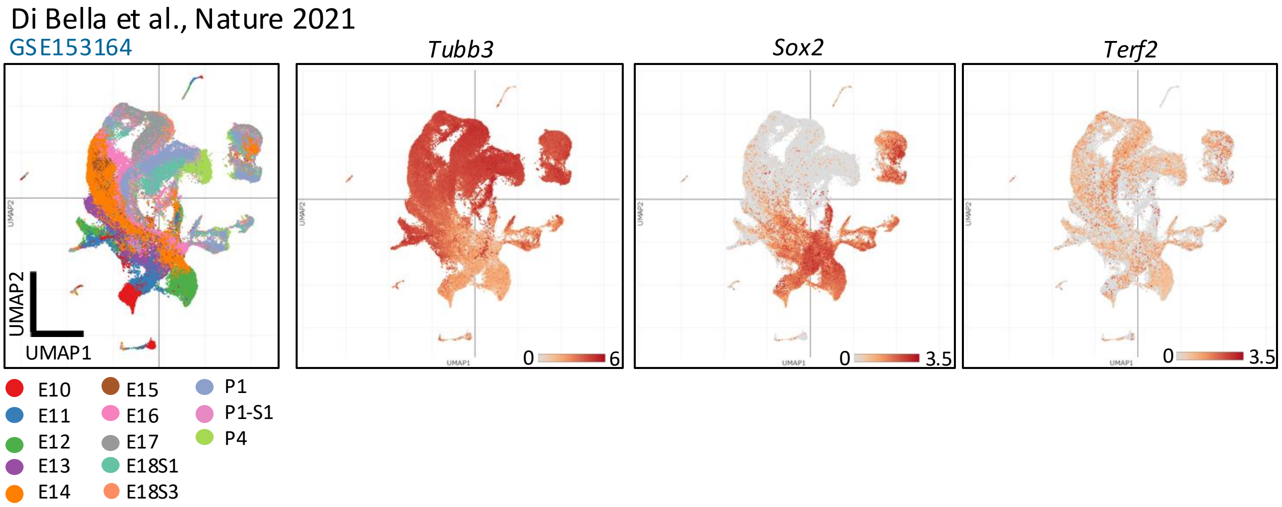


b

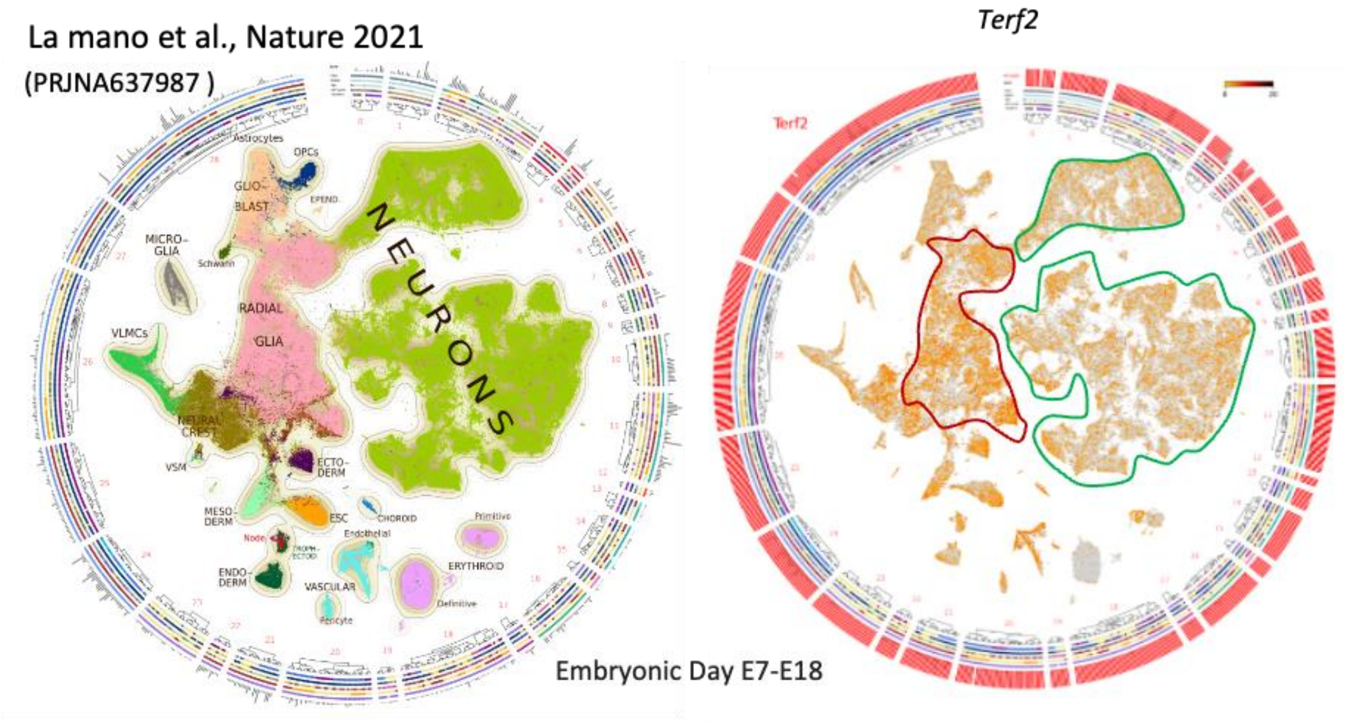

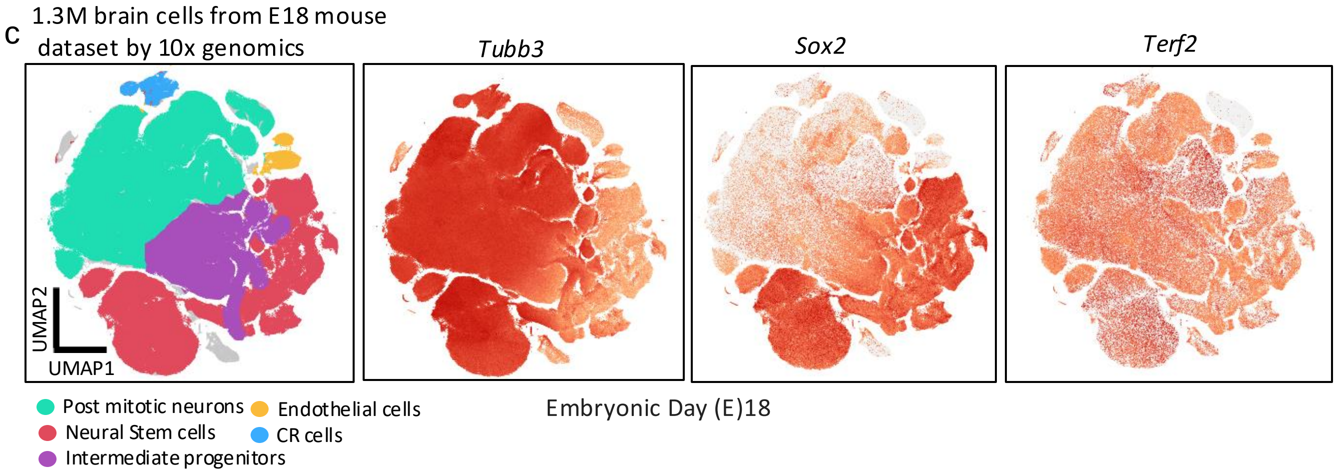


(A) Open access single cell RNA-seq data of embryonic mouse brain (E10-P4) highlights the presence of *Terf2* in both NSC (SOX2+ve) and Neuronal clusters (Tubb3+ve) (https://singlecell.broadinstitute.org/single_cell/study/SCP1290/molecular-logic-of-cellular-diversification-in-the-mammalian-cerebral-cortex#study-summary.)

(B) Wheel plot from the Atlas of developing mouse brain (http://mousebrain.org/wheel/) depicts expression of Terf2 in both Radial glia and neuronal clusters.

(C) Publicly available Single cell RNA seq data from E18.5 embryonic mouse brain (https://www.10xgenomics.com/datasets/1-3-million-brain-cells-from-e-18-mice-2-standard-1-2-0) shows expression of Terf2 in both NSC (SOX2+ve) and Neuronal clusters (Tubb3+ve).

**Supplementary Figure S1.4**

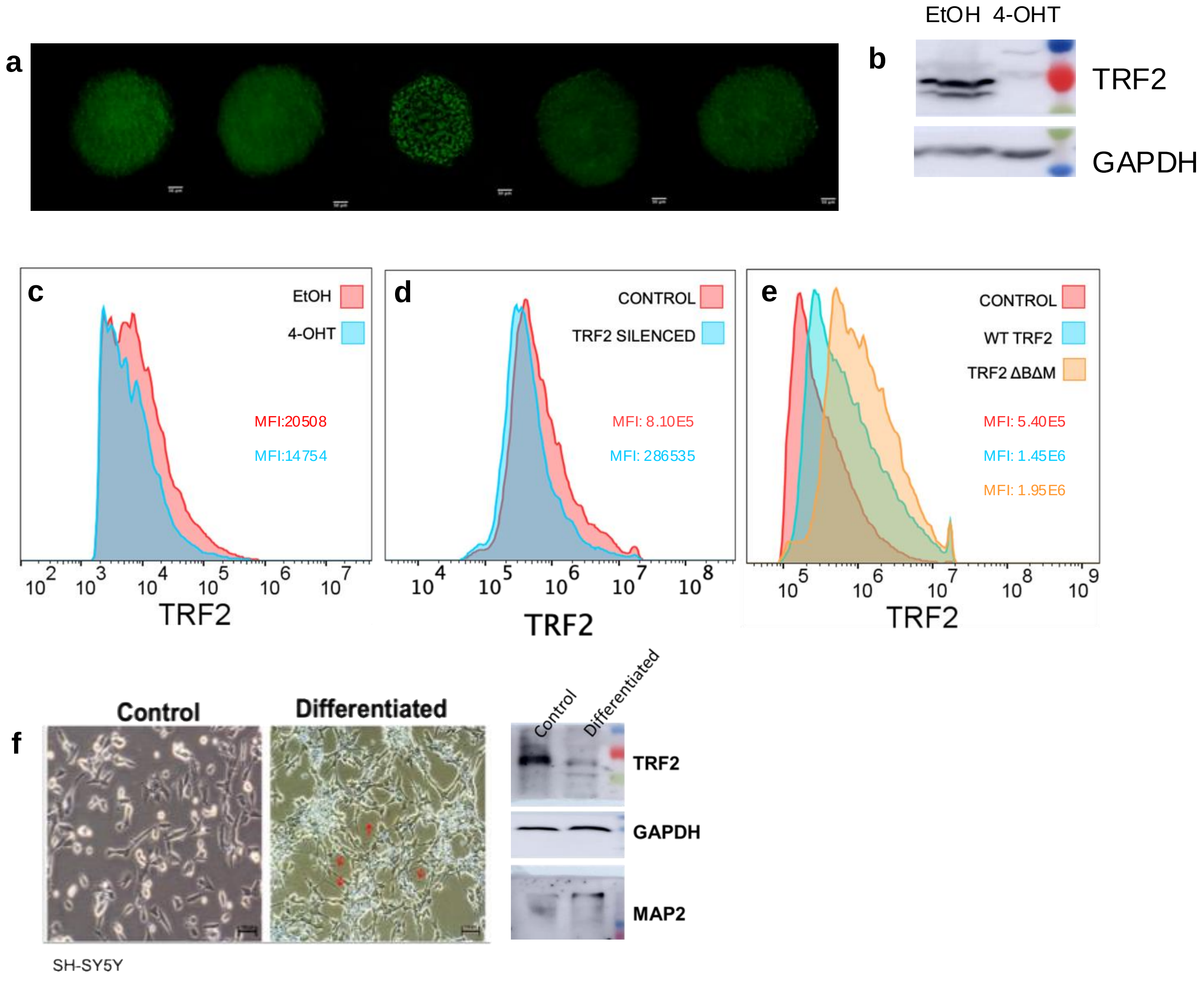


a. Immunofluorescence staining for PAX6 neurospheres from *Terf2$^{F/F}$/Nes:Cre* mice, stained using Alexa fluor-488 (green signal). Scale (50 µm)

b. TRF2 levels upon 4-OHT treatment in neurospheres.

c. Flow cytometry of mNSCs from *Terf2$^{F/F}$/Nes:Cre* mice; Vehicle control (EtOH) and TRF2 knockout cells (4-OHT). Representative Mean intensity of fluorescence (MIF) for TRF2 is shown. The cell counts are normalized to respective modes for comparative representation.

d. Flow cytometry of Control and TRF2 silenced SH-SY5Y cells. Representative Mean intensity of fluorescence (MIF) for TRF2 (supplementary) is shown. The cell counts are normalized to respective modes for comparative representation.

e. Flow cytometry of Control , WT TRF2 and TRF2 ΔBΔM. Representative Mean intensity of fluorescence (MIF) for TRF2 (supplementary) is shown. The cell counts are normalized to respective modes for comparative representation.

f. TRF2 levels post neuronal differentiation of SH-SY5Y cells.

**Supplementary Figure (for Figure 2)**

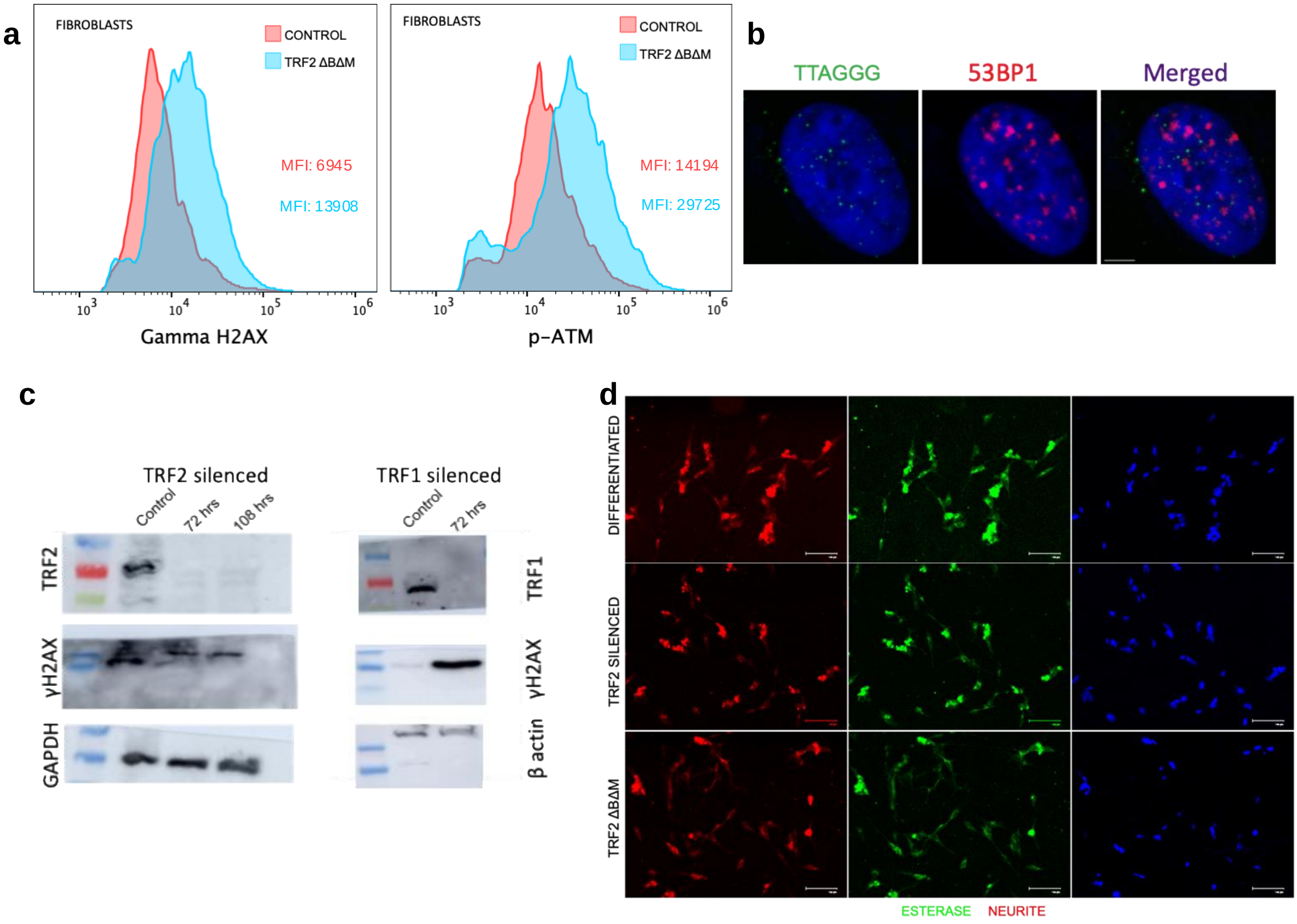


a. Flow cytometry using dual staining for γH2AX (i), p-ATM (ii) in SH-SY5Y cells and Fibroblasts. Mean intensity of fluorescence (MIF) for for γH2AX (i), p-ATM (ii) is shown. The cell counts are normalized to respective modes for comparative representation

b. (IF–FISH) for 53BP1 (red) and telomeres (green) in SHSY5Y treated with doxorubicin.

c. Western blot for  γH2AX in TRF2 and TRF1 silenced conditions.

d. Immunofluorescence staining for esterase and neurite in differentiated,TRF2 silenced and TRF2 ΔBΔM overexpression conditions. Esterase and Neurite were stained using neurite outgrowth staining kit 488 (red signal) and 594 (green signal), respectively. Scale bar (100 µm).

**Supplementary Figure (for fig 3)**.

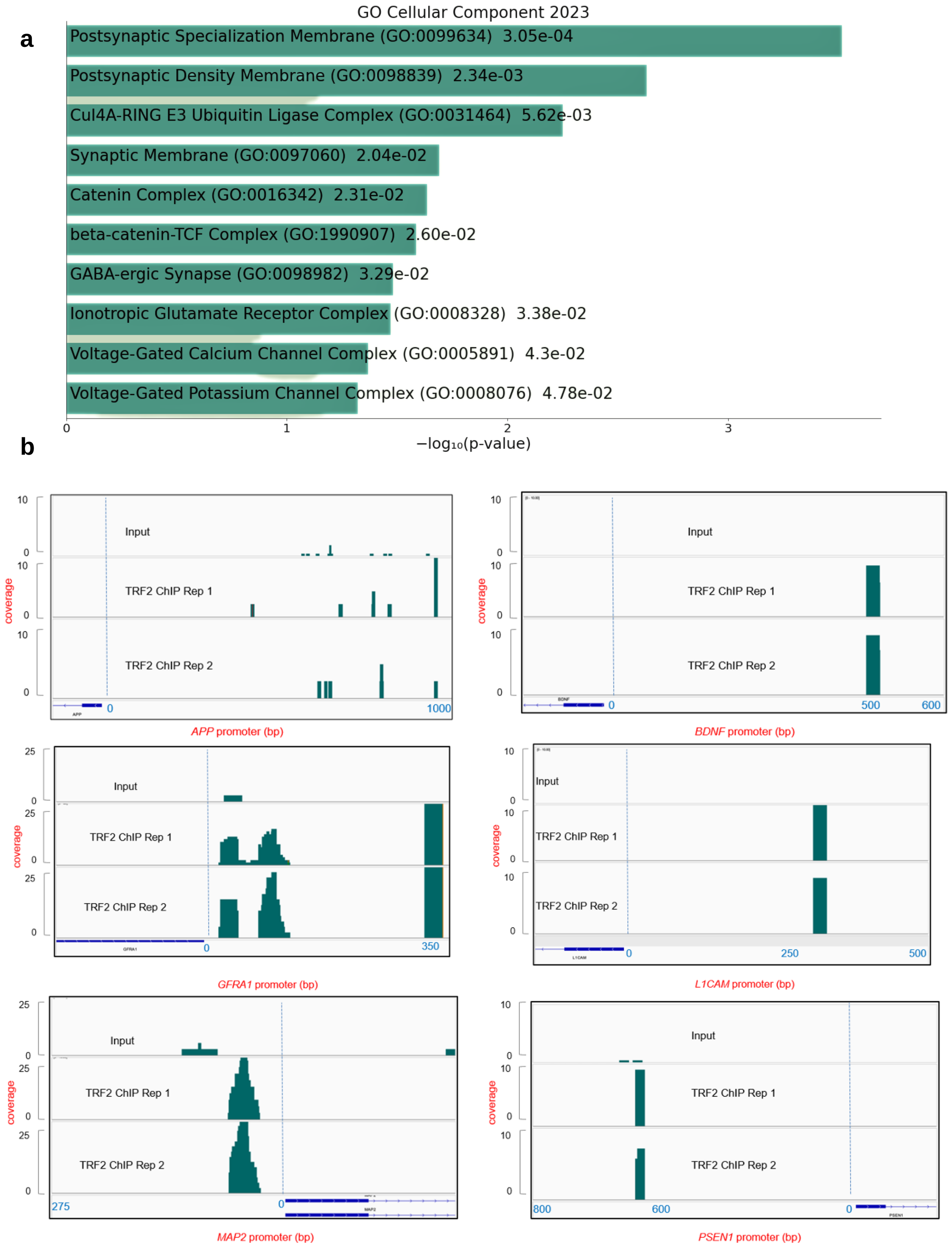


a. Gene Ontology (GO) Enrichment Analysis Using the ENrichr Database: The top 10 enriched terms for the TRF2 ChIP-seq peak annotated gene set are displayed based on the -log10(p-value), with the actual p-value shown next to each term.

b. Reads for TRF2 ChIP seq peak on the selected promoters of differentiation genes from two biological replicates in HT1080 cells.

**c**

| Human | Pimers used for ChIP- qRT PCR | Region covered upstream TSS |
|---|---|---|
| APP F | GACTCTCCCTCCCACTGTTC | |
| APP R | CGAGAGAGACCCCTAGCGG | 60-250 |
| BDNF F | CGAGATTTCCGGAGTCGT | |
| BDNF R | TGGAGGCGAAAGGATGAACT | 60-250 |
| GFRA1 F | CACATGCACCCCGACCTG | |
| GFRA1 R | GCTTTGAGATGAGAGCGGAG | 50-200 |
| L1CAM F | AGGGATACCCTCTCCCAGAC | |
| L1CAM R | CCCAGTCAGGGTCCTGTT | 125-400 |
| MAP2 F | CTGTGGTTGGGTTTGTGAGG | |
| MAP2 R | CACTGGCGTCCTTGGAAAAT | 60-250 |
| PSEN1 F | CTTCCTCCTGGCTCCTCC | |
| PSEN1 R | AGCTCAGGTTCCTTCCAGAC | 0-150 |

| Mouse | Pimers used for ChIP- qRT PCR | Region covered upstream TSS |
|---|---|---|
| APP F | CAAGACGCCGCACCTGTG | |
| APP R | AGAGTCAGCTGATCCCGC | 15-170 |
| BDNF F | GTCGTCCCCTTTAAGCAGC | |
| BDNF R | CACCCAGAGCTCAGCCAG | 10-160 |
| GFRA1 F | TAGTGATCGAGCCAGCAAGT | |
| GFRA1 R | ACCATTACTTTCAGAGCAAGCG | 10-180 |
| L1CAM F | AGTGGAGGCGAGGTAATCTG | |
| L1CAM R | CTCCAGCAGTCCCAGTATCC | 75-230 |
| MAP2 F | GGAAGGAAGGAGCCAAGTCT | |
| MAP2 R | GTCGCAGTGTAGGGGAGG | 50-200 |
| PSEN1 F | GTCGCGCACATCCTTACATC | |
| PSEN1 R | GACGTTGGCCCCTAGACC | 20-150 |

**Supplementary Figure (Fig 4)**.

a

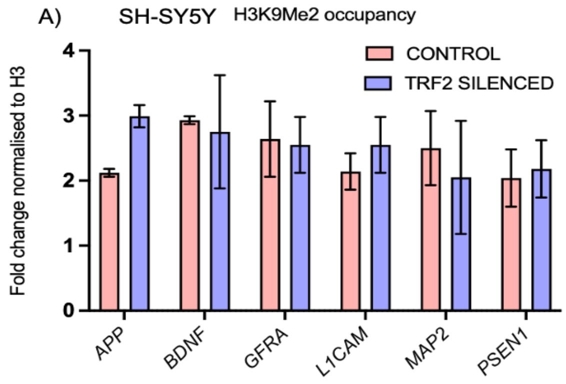


b

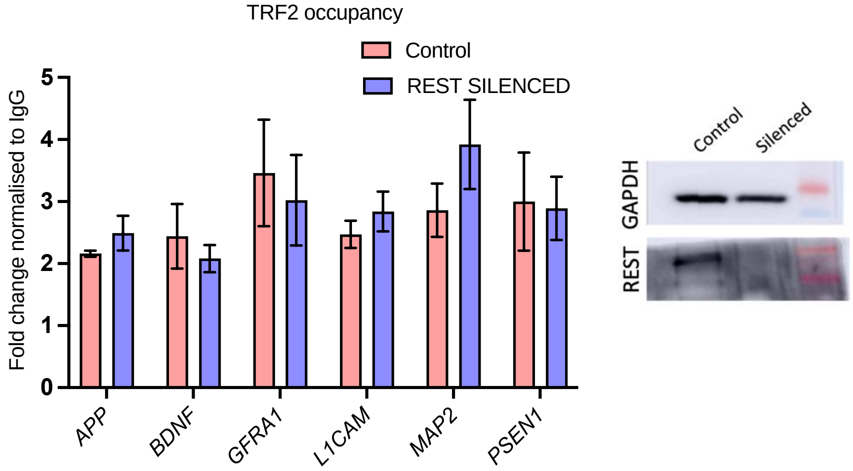


c

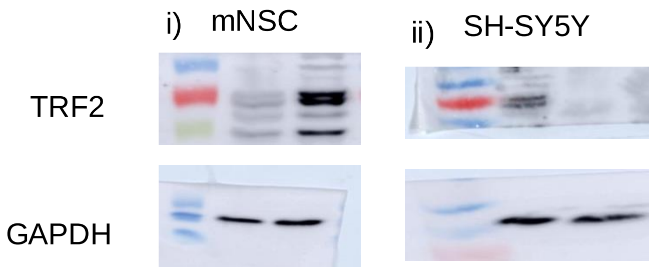


a. H3K9Me2 occupancy upon TRF2 silencing in SHSY5Y cells.

b. REST occupancy upon TRF2 silencing in SH-SY5Y cells.

c. Representatiive silencing levels of TRF2 in i) mNSC and ii) SH-SY5Y

**Supplementary Figure (for fig 5)**

**a**

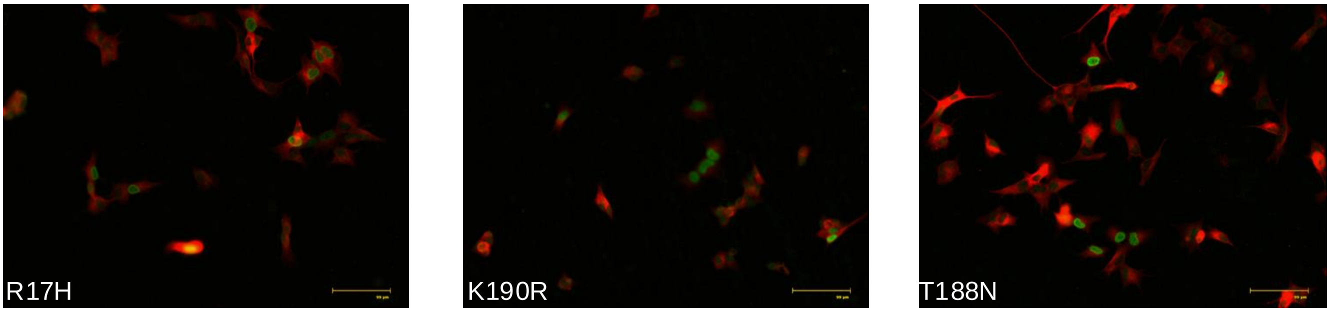


**b**

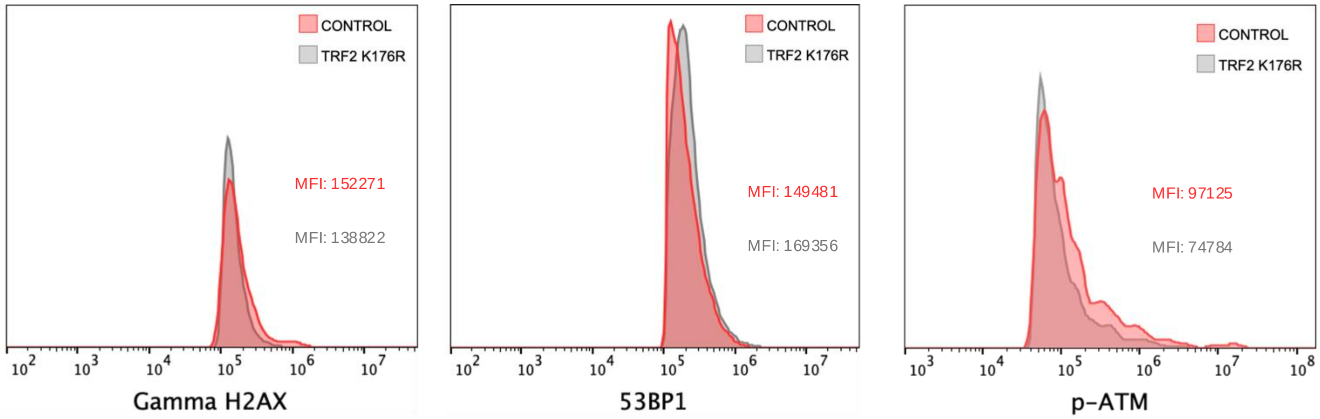


a. Immunofluorescence staining for TRF2 and β- III Tubulin proteins in TRF2 R17H, K90R and T188N overexpression conditions. β- III Tubulin and TRF2 were stained using Alexa fluor-594 (red signal) and Alexa fluor-498 (green signal), respectively.

b. Flow cytometry with staining for γH2AX (i), 53BP1 (ii), p-ATM (iii) in SH-SY5Y cells. Control and TRF2 K176R overexpression cells. Mean intensity of fluorescence (MIF) for γH2AX (i), 53BP1 (ii), p-ATM (iii). The cell counts are normalized to respective modes for comparative representation.

**Supplementary Figure (for fig 6)**

a. Promoter sequences of differentiation genes containing G-quadruplex (G4) motifs were analyzed. Panel A presents a table detailing the location, distance from the transcription start site (TSS), and the G4 motifs, including both wild-type sequences and their corresponding G4 mutant variants, within promoter regions that coincide with TRF2 promoter binding.

b. Circular dichroism (CD) spectra of the PG4-motif sequences presented in the table above were analyzed, with base-substituted mutant sequences that are not capable of forming G-quadruplex (G4) structures serving as controls.

c. Structure of ligands SMH14.6 and Bis indole carboxamide.

**a**

| Sequence name | Sequence | Distance from TSS |
|---|---|---|
| L1CAM CD _H_G3 | GGGCGGGGCGGGCGGCCGGG | 125 |
| L1CAM Mut_H_G3 | GTGCGTGGCGTGCGGCCGTG | |
| MAP2 CD _H | GGGACCTGCGCGGGGCGGGGAGGAGGG | 48 |
| MAP2 Mut_H | GTGACCTGCGCGTGGCGGTGAGGAGTG | |
| APP_ CD _H | GGGTCTCTCTCGGGTGCCGAGCGGGGTGGG | 57 |
| APP_ Mut_H | GTGTCTCTCTCGTGTGCCGAGCGGGGTGGG | |
| BDNF_ CD _H | GGGGGGCGGGGGGCGGGGGGGGGGGGG | 66 |
| BDNF_Mut_H | GTGTGTCTGGTGTCGTGGTGGTGTGTT | |
| GFRA_ CD _H | GGGAACGGGGGAGGGGAGAGGG | 112 |
| GFRA_Mut_H | GTGAACGTGGGAGTGGAGAGTG | |
| PSEN1_CD _H | GGGTTCTCGGGCGGGGCCTGGG | 33 |
| PSEN1_ Mut_H | GTGTTCTCGTGCGGTGCCTGTG | |
| L1CAM CD _M | GGCCTGCTGCGGGGGAGG | 82 |
| L1CAM Mut_M | GGCCTGCTGCGTGGGAGT | |
| MAP2 CD _M | GGCGGCGCTCGGGCTGCGCGG | 56 |
| MAP2 Mut_M | GGCGTCGCTCGGGCTGCGCGT | |
| APP_ CD _M | GGGTCTCTGTCGGGTGGCGAGCCGGGTGGG | 53 |
| APP_ Mut_M | GTGTCTCTGTCGTGTGGCGAGCCGTGTGTG | |
| BDNF_ CD _M | GGAGCAGGGAGGGGATAGGG | 68 |
| BDNF_Mut_M | GTAGCATGGAGGGTATAGGT | |
| GFRA_ CD _M | GGGAGCGGGGGAGGGGAGAGGG | 22 |
| GFRA_Mut_M | GTGAGCGTGGGAGTGGAGAGTG | |
| PSEN1_CD _M | GGCGGGGCCTAGG | 13 |
| PSEN1_ Mut_M | GTCGTGGCCTAGG | |

**b**

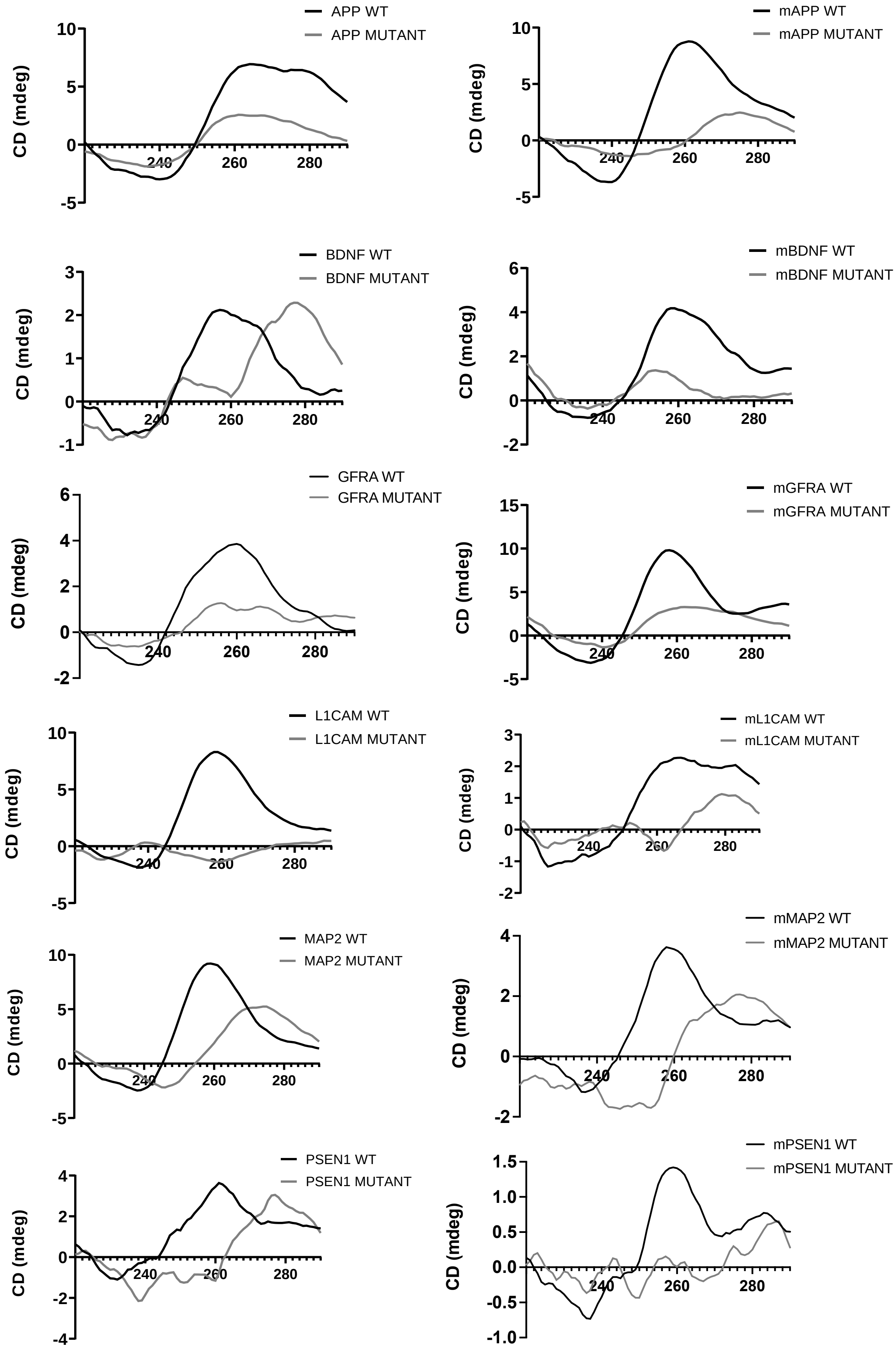
APP WT
APP MUTANT
mAPP WT
mAPP MUTANT
BDNF WT
BDNF MUTANT
mBDNF WT
mBDNF MUTANT
GFRA WT
GFRA MUTANT
mGFRA WT
mGFRA MUTANT
L1CAM WT
L1CAM MUTANT
mL1CAM WT
mL1CAM MUTANT
MAP2 WT
MAP2 MUTANT
mMAP2 WT
mMAP2 MUTANT
PSEN1 WT
PSEN1 MUTANT
mPSEN1 WT
mPSEN1 MUTANT
CD (mdeg)
240
260
280

**c**

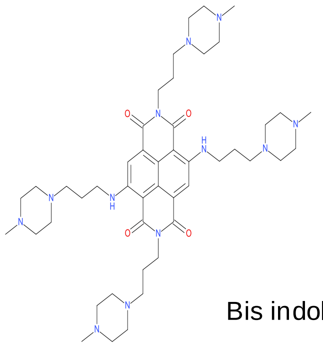

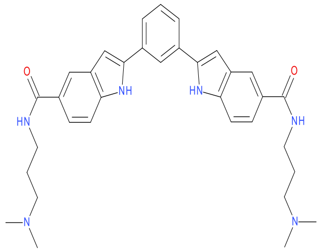

SMH14.6 Bis indole carboxamide
